# Transcriptional control of nucleus accumbens neuronal excitability by Retinoid X Receptor Alpha tunes sensitivity to drug rewards

**DOI:** 10.1101/2022.03.28.486101

**Authors:** Arthur Godino, Marine Salery, Romain Durand-de Cuttoli, Molly S. Estill, Leanne M. Holt, Rita Futamura, Caleb J. Browne, Philipp Mews, Peter J. Hamilton, Rachael L. Neve, Li Shen, Scott J. Russo, Eric J. Nestler

## Abstract

The complex nature of the transcriptional networks underlying addictive behaviors suggests intricate cooperation between diverse gene regulation mechanisms that go beyond canonical activity-dependent pathways. Here we implicate in this process a novel nuclear receptor transcription factor, Retinoid X Receptor Alpha (RXRα), which we identified bioinformatically as associated with addiction-like behaviors. In the nucleus accumbens (NAc) of male and female mice, we show that, while its own expression remains unaltered after cocaine exposure, RXRα controls plasticity- and addiction-relevant transcriptional programs in both dopamine receptor D1- and D2-expressing medium spiny neurons, which in turn modulate intrinsic excitability and synaptic activity of these NAc cell types. Behaviorally, bidirectional viral and pharmacological manipulation of RXRα regulates drug reward sensitivity in both non-operant and operant paradigms. Together, this study demonstrates a key role for NAc RXRα in promoting drug addiction, and paves the way for future studies of rexinoid signaling in psychiatric disease states.

## INTRODUCTION

Drugs of abuse such as cocaine perturb coordinated activity within the brain’s reward circuitry ^1^ initially in part by increasing dopamine signals to levels far exceeding those of natural reinforcers ^2^, thus hijacking more classic mechanisms of reward learning ^3^ to build pathological drug memories upon repeated exposure ^4^. The transition to a compulsive state of drug seeking and taking even in spite of negative consequences – a behavioral hallmark of substance use disorder ^5^ – however results from a composite interplay between a drug’s pharmacological properties and an individual’s sensitivity to drug reward, which itself depend on both innate and environmental factors ^6–9^.

Genetic, pharmacological and environmental effects converge onto the establishment of drug-induced transcriptional programs that underlie the several molecular-, cellular-, synaptic-, circuit- and behavioral-level alterations that define the drug-addicted phenotype ^4, 10^. In a recent effort to better link brain-wide transcriptional patterns with individual behavioral responses to cocaine, we performed RNA-sequencing (RNAseq) on six regions of the mouse brain’s reward circuitry after cocaine self-administration, withdrawal, and relapse (Fig. 1a) ^11^. One major finding of this study was the large differences in transcriptional landscapes across time points and brain regions ^11^. This indicated that the sole recruitment of canonical activity-dependent signaling pathways upstream of transcription factors such as CREB or ΔFosB – previously extensively studied in addiction ^12^ – could not account fully for such heterogeneity in transcriptional responses. Subsequent *in silico* analyses ^11^ led us to propose that nuclear receptors – a large but understudied family of ligand-activated transcription factors that have the ability to form heterogeneous dimers and thus to coordinate gene expression across pathways yet with high specificity ^13^ – could be critical co-regulators of temporal- and region-specific transcriptional programs.

**Fig. 1.**
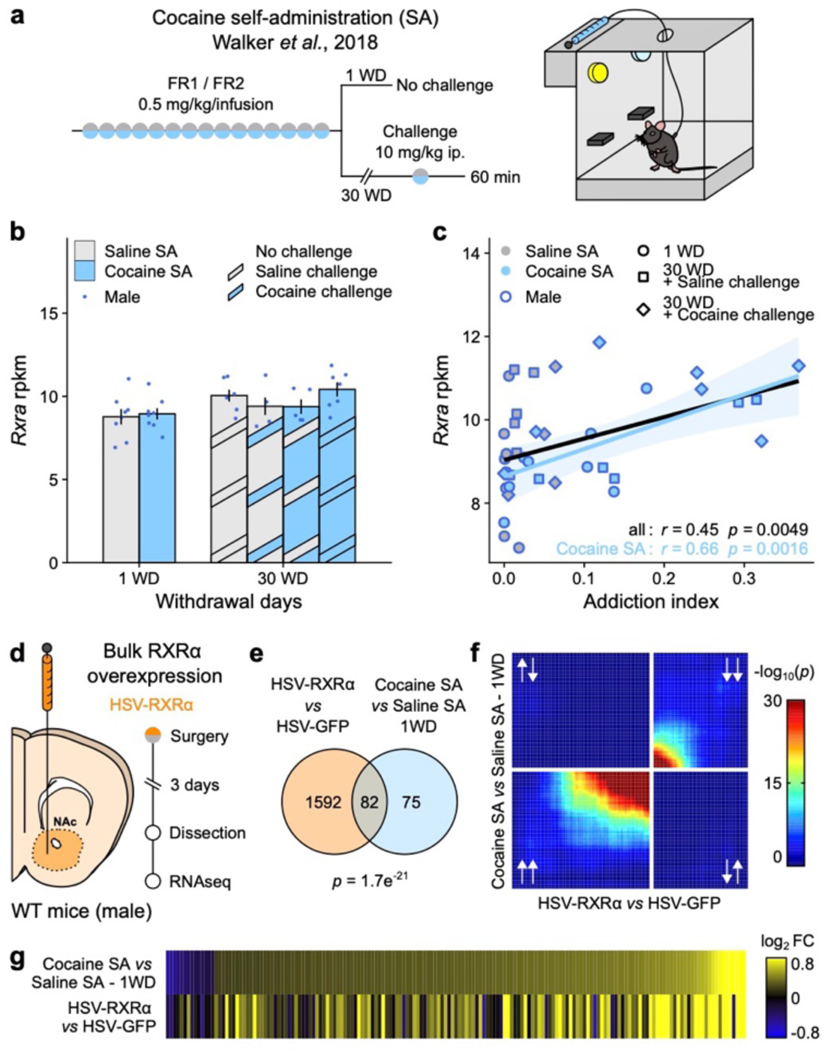
RXRα expression in NAc is linked to addiction-relevant behavior and transcription. **a**, Experimental design of the original cocaine self-administration (SA) study ^11^. Male mice (n = 5-8/group) were subjected to 10-15 days of cocaine SA and euthanized after either 1 or 30 days of withdrawal (WD) in their home cage. Mice in the 30 WD group were given a challenge injection of cocaine or saline and re-exposed to their original SA chamber for 1 hour before being euthanized. **b**, RNAseq of NAc tissue showed no regulation of *Rxra* transcripts across experimental groups (data from original study ^11^). **c**, *Rxra* levels in NAc correlated with the Addiction Index, a composite metric representative of addiction-like behavioral domains computed using exploratory factor analysis ^11^ (across all samples: Pearson’s *r* = 0.45, *p* = 0.0049; across cocaine SA samples only: Pearson’s *r* = 0.66, *p* = 0.0016). **d**, Experimental design of bulk RXRα overexpression transcriptomics study. Male mice (n = 4/group) were injected in NAc with an HSV construct overexpressing RXRα or a control GFP sequence, then euthanized 4 days later. **e**, RNAseq of virally-infected NAc tissue revealed a significant overlap between transcripts significantly regulated by RXRα overexpression and following 1 WD from cocaine SA ^11^ (Fisher’s exact test *p* = 1.7e^−21^). **f**, Comparison of transcriptome-wide expression profiles using rank-rank hypergeometric overlap (RRHO) plots confirmed overlap between the two datasets in a threshold-free manner (lower-left quadrant: genes upregulated in both studies, upper-left quadrant: genes upregulated by RXRα overexpression but downregulated by Cocaine S1 + 1 WD, lower-right quadrant: genes downregulated by RXRα overexpression but upregulated by Cocaine S1 + 1 WD, upper-right quadrant: genes downregulated in both studies). **g**, Similar patterns of gene expression visualized as heatmaps of expression fold changes from respective controls. Bar graphs represent mean ± sem. Correlation graphs represent regression line with its 95% confidence interval.

In the nucleus accumbens (NAc), a key striatal region of the brain’s reward circuitry that integrates midbrain dopaminergic inputs together with cortical and limbic glutamatergic signals to shape reward- and motivation-related behaviors ^14^, the nuclear receptor Retinoic X Receptor Alpha (RXRα) stood out. Its transcript levels were not altered by cocaine exposure (Fig. 1b) ^11^ but positively correlated with the “Addiction Index”, a bioinformatically-derived composite metric of addiction-relevant behaviors (Fig. 1c) ^11^, altogether making RXRα a particularly suitable mechanistic candidate to explain individual vulnerability to cocaine addiction. While burgeoning evidence implicates retinoic acid-related signaling ^15–17^ and other RXR family members – especially RXR Gamma (RXRγ) ^18–21^ – in striatal-dependent motivation-related behaviors, data on RXRα in the mature brain remains scarce. Genome-wide association studies have linked human *Rxra* mutations with schizophrenia ^22–24^, cognitive ability ^25, 26^ and dementia ^27, 28^, and a recent study demonstrated that RXRα can control spine and synapse formation in the adult mouse cortex ^29^, suggesting a role for RXRα in brain plasticity mechanisms.

Therefore, we hypothesized RXRα to be an important transcriptional regulator of NAc function in drug-related behaviors. One key question was also to identify whether RXRα action in NAc would be cell-type-specific. Indeed, the NAc is composed mainly of two largely non-overlapping subpopulations of GABAergic medium spiny neurons (MSNs) that express either the dopamine D1 (*Drd1*) or dopamine D2 receptor (*Drd2*) ^30^, and which have been shown to play different – even antagonistic – roles in drug-related motivated and reward-learning behaviors ^31–33^. To that end, we here assess RXRα regulation after drug exposure, as well as the transcriptional, physiological and behavioral consequences of viral manipulation of RXRα levels in D1- and D2-MSNs to propose a model for RXRα‘s contribution to drug reward and addiction. We also provide early preclinical evidence for targeting RXRα using small molecule inhibitors as a possible new pharmacotherapeutic avenue for patients with substance use disorder.

## RESULTS

### RXRα mediates addiction-like transcriptional programs

In a previous brain-wide transcriptomics study following cocaine self-administration (Fig. 1a) ^11^, *Rxra* transcript levels were not affected at any of the time points analyzed in the NAc (Fig. 1b) or in any of the five other brain regions analyzed in that study (Extended Data Fig. S1a) ^11^. However, correlation between individual *Rxra* transcript levels and the Addiction Index, a multi-factorial metric summarizing addiction-like behavioral features, was the strongest in NAc (Fig. 1c and Extended Data Fig. S1b, Pearson’s *r* = 0.45, *p* = 0.0049), even when only considering cocaine-exposed animals (Pearson’s *r* = 0.66, *p* = 0.0016), ruling out a Simpson’s paradox effect. This analysis suggests that pre-existing individual levels of RXRα expression in NAc might influence the severity of addiction-like behaviors upon drug exposure, presumably by contributing to cocaine-induced gene regulation mechanisms in this brain region. To test this hypothesis, we compared genes regulated in NAc by RXRα to those regulated by cocaine exposure in this original study ^11^. In order to detect direct or indirect gene targets of RXRα, we performed bulk RNAseq on RXRα-overexpressing or control NAc tissue (Fig. 1d). RXRα overexpression was achieved by infusing a Herpes Simplex Virus (HSV) encoding RXRα (or GFP as a control) and confirmed at both the RNA and protein levels (Extended Data Fig. S2). Among the 1674 genes significantly regulated by RXRα overexpression (all genes and corresponding statistics available in Extended Data Table S1), 82 were also regulated in NAc one day after cocaine self-administration – more than 50% of the 157 genes affected in that condition (Fig. 1e). This significant overlap (Fisher’s exact test *p* = 1.7e^−21^) strengthened the idea of RXRα contributing to NAc transcriptional programs in response to chronic self-administered cocaine. Subsequent comparisons using rank-rank hypergeometric overlap (RRHO) plots (Fig. 1f) or expression-based heatmaps (Fig.1g) highlighted overlapping genes as being mostly upregulated, suggesting that RXRα might predominantly act as a permissive transcription factor ^13, 34, 35^ in that context. Collectively, these initial findings warranted further study of RXRα in the transcriptional mechanisms of cocaine reinforcement.

### Acute cocaine does not affect RXRα expression

Our next goal was to examine RXRα regulation at both RNA and protein levels. Male and female mice were injected intraperitoneally with a 20 mg/kg dose of cocaine, and NAc tissue dissected 30 or 60 min after the injection (Fig. 2a). We first confirmed the absence of regulation of *Rxra* transcript levels after acute cocaine in males and females (Fig. 2b, statistics in Extended Data Table S6). In addition, no significant changes in RXRα expression were detected at the total protein level (Fig. 2c, statistics in Extended Data Table S6). Another process through which RXRα could mediate its genomic effects is through regulation of its nuclear localization ^29^. Here, we did not observe any changes in RXRα protein levels in nuclear-enriched fractions from NAc tissue after acute cocaine treatment (Fig. 2d, statistics in Extended Data Table S6). However, this does not exclude regulation via other mechanisms like phosphorylation or truncation ^34^, or regulation in other more complex experimental settings.

**Fig. 2.**
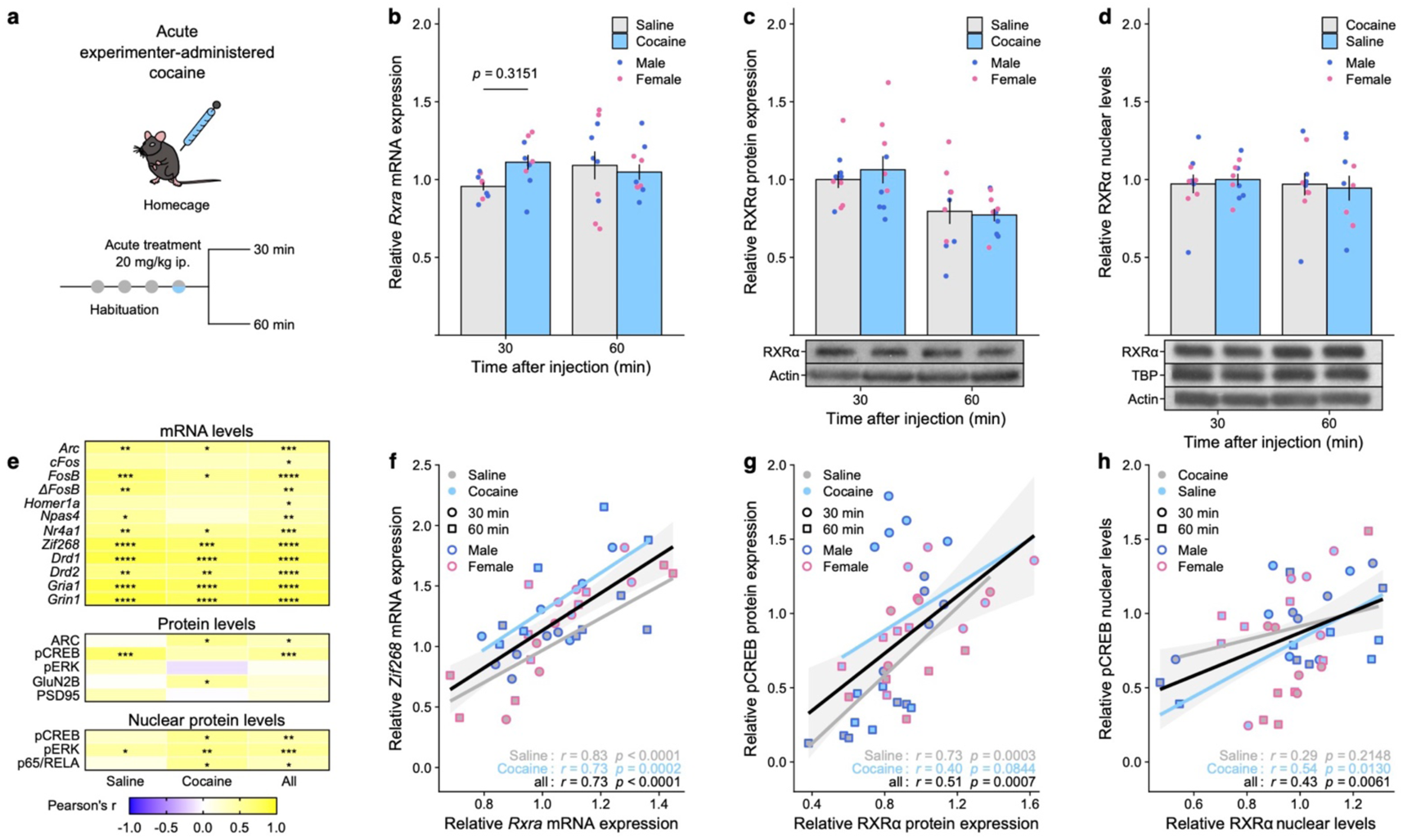
RXRα expression levels in NAc are not regulated by acute cocaine but correlate with markers of striatal activity and plasticity. **a**, Experimental design. Following habituation, male and female mice (n = 4-5/group) were injected with an acute dose of cocaine (20 mg/kg) or saline and euthanized 30 or 60 min later for dissection of NAc tissue. **b**, *Rxra* mRNA levels were not significantly regulated after acute cocaine exposure (LMM-ANOVA: main effect of Drug: F_1,31_ = 0.8185, *p* = 0.3726; interaction Drug:Time: F_1,31_ = 2.5762, *p* = 0.1186, followed by Sidak’s *post hoc* tests). **b**, RXRα protein levels in whole tissue lysates were not significantly regulated after acute cocaine exposure (LMM-ANOVA: main effect of Drug: F_1,32_ = 0.1004, *p* = 0.7534; interaction Drug:Time: F_1,32_ = 0.4864, *p* = 0.4906, followed by Sidak’s *post hoc* tests). **c**, RXRα protein levels in nuclear fractions were not significantly regulated after acute cocaine exposure (LMM-ANOVA: main effect of Drug: F_1,32_ = 0.4631, *p* = 0.2353; interaction Drug:Time: F_1,32_ = 0.0070, *p* = 0.9340, followed by Sidak’s *post hoc* tests). **e,** Summary heatmap showing that *Rxra*/RXRα levels correlated strongly with other genes/proteins implicated in striatal function – including several IEGs – across saline only, cocaine only or all samples (Pearson’s *r*, * *p* < 0.05, ** *p* < 0.01, *** *p* < 0.001, **** *p* < 0.0001). **f**, Representative example of *Rxra* mRNA levels correlation with the IEG *Zif268* (across saline samples only: Pearson’s *r* = 0.83, *p* < 0.0001; across cocaine samples only: Pearson’s *r* = 0.73, *p* = 0.0002; across all samples: Pearson’s *r* = 0.73, *p* < 0.0001). **g**, Representative example of RXRα protein levels correlation with active transcription factor pCREB (across saline samples only: Pearson’s *r* = 0.73, *p* = 0.0003; across cocaine samples only: Pearson’s *r* = 0.40, *p* = 0.0844; across all samples: Pearson’s *r* = 0.51, *p* = 0.0007). **h**, Representative example of RXRα nuclear levels correlation with pCREB (across saline samples only: Pearson’s *r* = 0.29, *p* = 0.2148; across cocaine samples only: Pearson’s *r* = 0.54, *p* = 0.0130; across all samples: Pearson’s *r* = 0.43, *p* = 0.0061). Representative Western Blot pictures for RXRα and control actin and TBP bands are attached to the corresponding quantification. Bar graphs represent mean ± sem after combining male and female data. Correlation graphs represent regression line with its 95% confidence interval.

### RXRα expression levels correlate with markers of striatal function

Following up on the original conjecture that NAc RXRα levels might control drug-evoked behavioral responses, we checked whether RXRα levels would also predict drug-evoked molecular responses by correlating individual RXRα transcript, total protein and nuclear levels with markers of striatal function and drug-induced plasticity. We found striking positive correlations with expression levels of several key players, including dopamine (*Drd1*, *Drd2*) and glutamate (*Gria1*, *Grin1*, GluN2B) receptors, as well as drug-related signaling molecules like phosphorylated ERK and NFκB-complex transcription factor p65/RELA (Fig. 2e, statistics in Extended Data Table S6). We also found significant positive correlations between RXRα and expression levels of all immediate early genes (IEGs) tested, including *Zif268* (Fig. 2f), which are well-established proxies of stimulus-induced neuronal activation and key intermediates in drug-induced molecular plasticity ^36^. Of note, a causal link between RXRα action and IEG expression has been proposed before ^29^. At the protein level in both total (Fig. 2g) and nuclear (Fig. 2h) extracts, RXRα positively correlated with levels of phosphorylated (active) CREB (pCREB), the convergence point of many drug-triggered signaling cascades ^12^. Although not a causal mechanistic explanation, these correlative data provide first clues as to the molecular underpinnings of RXRα striatal function: RXRα might control select target transcriptional programs that affect MSN excitability and activation of stimulus-induced signaling pathways, such as – but likely not limited to – pCREB-mediated IEG induction.

### RXRα is partly enriched in NAc D1-MSNs

A growing literature underscores the importance of considering cell-type specificity in studies of drug-induced signaling and plasticity ^12, 33^. Accordingly, we checked for RXRα enrichment in the two main NAc cell types, which express either *Drd1* or *Drd2* ^30^ using RNA fluorescent *in situ* hybridization (FISH; Fig. 3a). Quantification of *Drd1* and *Drd2* probes in DAPI-identified nuclei allowed for robust classification into D1-MSNs (*Drd1*^+^/*Drd2*^−^) or D2-MSNs (*Drd1*^−^/*Drd2*^+^) (Extended Data Fig. S3). Quantification of *Rxra* puncta within cell-type-classified nuclei revealed a significant 18 % higher *Rxra* expression level in D1-MSNs compared to D2-MSNs (LMM-ANOVA: main effect of CellType: F_3,19381_ = 45.1531, *p* < 2.2e^−16^, followed by Sidak’s *post hoc* tests).

**Fig. 3.**
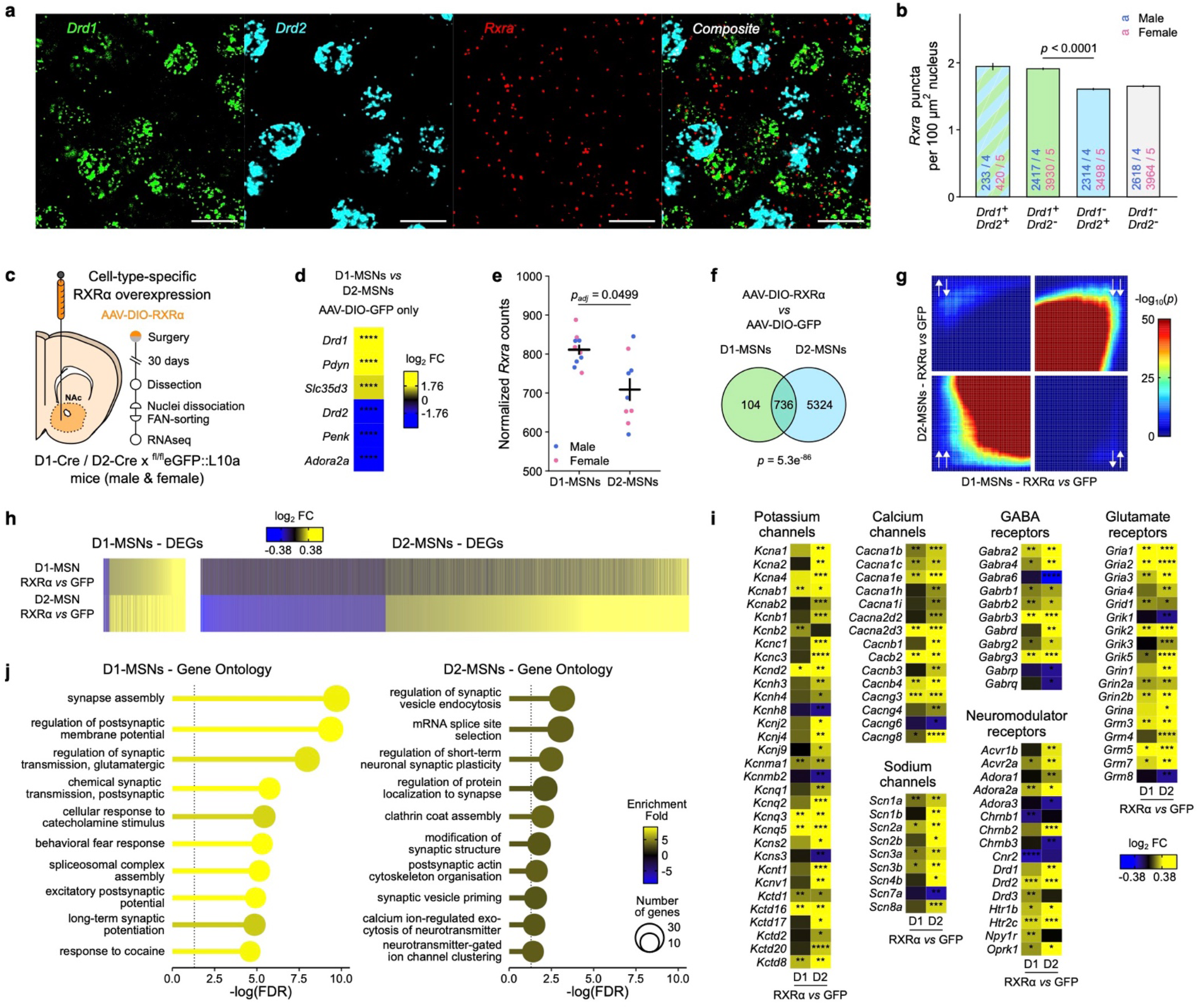
RXRα controls plasticity- and excitability-related transcriptional programs in both D1- and D2-MSNs. **a**, Representative images of RNA FISH for *Drd1*, *Drd2* and *Rxra* mRNAs in NAc. Scale bar is 20 µm. **b**, Quantification of *Rxra* mRNA puncta in nuclei of NAc cell types classified on *Drd1* and *Drd2* expression levels in both male and female mice (n = 4-5/group) showed higher *Rxra* expression in D1-MSNs at baseline. (LMM-ANOVA: main effect of CellType: F_3,19381_ = 45.1531, *p* < 2.2e^−16^, followed by Sidak’s *post hoc* tests). **c**, Experimental design of cell-type-specific RXRα overexpression transcriptomics study. Male and female mice (n = 4-5/group) from D1-Cre x ^fl/fl^eGFP::L10a or D2-Cre x ^fl/fl^eGFP::L10a crosses were injected in NAc with an AAV construct overexpressing RXRα or a control GFP sequence in a Cre-dependent manner, then euthanized 30 days later. NAc tissue was dissected and dissociated into a single-nuclei suspension before FAN-sorting of GFP^+^ nuclei. **d**, RNAseq confirmed enrichment of D1-MSNs and D2-MSNs marker genes in sorted D1 and D2 nuclei, respectively (Wald’s test D1-MSNs *vs* D2-MSNs, **** *p* < 0.0001). **e**, RNAseq also confirmed increased *Rxra* expression in D1-MSNs at baseline (Wald’s test with Benjamini-Hochberg correction for multiple comparisons *p_adj_* = 0.0499). **f**, Significant overlap between transcripts significantly regulated by RXRα overexpression in D1-MSN and D2-MSN nuclei (Fisher’s exact test *p* = 5.3e^−86^). **g**, Comparison of D1- and D2-specific transcriptome-wide expression profiles using rank-rank hypergeometric overlap (RRHO) plots confirmed overlap (lower-left quadrant: genes upregulated in both cell types, upper-left quadrant: genes upregulated in D1-MSNs but downregulated in D2-MSNs, lower-right quadrant: genes downregulated in D1-MSNs but upregulated in D2-MSNs, upper-right quadrant: genes downregulated in both cell types). **h**, Similar patterns of gene expression visualized as heatmaps of expression fold changes from respective controls. **i**, Selected excitability- and plasticity-related genes are regulated by RXRα overexpression in D1- or D2-MSNs (Wald’s test, * *p* < 0.05, ** *p* < 0.01, *** *p* < 0.001, **** *p* < 0.0001). **j**, Gene ontology analyses of all genes significantly regulated by RXRα overexpression indicated enrichment of genes involved in neuronal and synaptic function in both D1- and D2-MSN. Bar graphs represent mean ± sem after combining male and female data.

### RXRα controls similar excitability- and plasticity-related transcriptional programs in D1- and D2-MSNs

RXRα enrichment in D1-MSNs, along with published work on those cell types ^31–33^, justified considering cell-type-specificity in our investigation of the targets downstream of RXRα signaling. For this reason, we used male and female D1-Cre and D2-Cre mice – which express Cre recombinase under the promoter of the *Drd1* or *Drd2* gene – in combination with a Cre-dependent adeno-associated virus expressing RXRα (AAV-DIO-RXRα) infused in the NAc to achieve population-specific RXRα overexpression (Fig. 3a). These D1-Cre and D2-Cre mice were also crossed with a ^fl/fl^eGFP::L10a reporter line that allows for Fluorescence-Activated Nuclei Sorting (FANS) of D1- and D2-positive nuclei ^37^ (Extended Data Fig. 4). RXRα overexpression was confirmed in sorted cell types at the transcript level and in whole NAc tissue at the RNA and protein levels (Extended Data Fig. S2). First, we validated enrichment of canonical D1- and D2-MSNs markers ^37, 38^ in the corresponding populations (Fig. 3d, statistics in Extended Data Table S6). We also confirmed *Rxra* enrichment in D1-MSNs (Fig. 3e, Wald’s test with Benjamini-Hochberg correction for multiple comparisons *p_adj_* = 0.0499), consistent with RNA FISH data (Fig. 3b). Next, we compared genes regulated by RXRα overexpression (AAV-DIO-RXRα *vs* AAV-DIO-GFP) separately in D1-MSNs or D2-MSNs (all genes and corresponding statistics available in Extended Data Table S2 and S3). While the total number of differentially expressed genes (DEGs) at the predefined significance threshold (± 15% change, nominal *p* < 0.05) was much higher in D2-MSNs than D1-MSNs (6060 D2-MSNs DEGs *vs* 840 D1-MSNs DEGs), DEGs lists overlapped considerably (Fisher’s exact test *p* = 5.3e^−86^) between the MSN subtypes, with almost 90% of the D1-DEGs also passing significance criteria in D2-MSNs (Fig. 3f). Visualization in RRHO plots also revealed strong overlap in threshold-free transcriptomic landscapes associated with RXRα overexpression in the two cell types, for both up- and down-regulated genes (Fig. 3g). Interestingly, expression fold-change heatmaps illustrated that, even for genes that do not reach significance, the change direction (up/down-regulation) was conserved across cell types (Fig. 3h), suggesting that RXRα regulates extremely similar gene programs, but that the magnitude of these changes is blunted in D1-MSNs relative to D2-MSNs. We examined DEGs that encode potassium, calcium and sodium channels, as well as GABA, glutamate or neuromodulator receptors, which recapitulated the same pattern of larger fold change magnitude in D2-MSNs (Fig. 3i, statistics in Extended Data Table S6). Unbiased gene ontology enrichment analyses further examined involvement of RXRa-regulated genes in neuronal excitability, signaling and plasticity mechanisms (Fig. 3j). Yet, enrichment for these excitability- or plasticity-associated gene ontology terms was much higher in D1-MSNs than D2-MSNs (Fig.3i). This finding suggests that RXRα-dependent transcriptional programs in D2-MSNs, although larger in number of DEGs, are less explicitly related to excitability or plasticity processes than in D1-MSNs. Despite these differences, this RNAseq dataset demonstrates that RXRα governs extremely similar transcriptional programs in D1- and D2-MSNs, which include numerous and diverse candidate effector genes that participate in neuronal physiology.

**Fig. 4.**
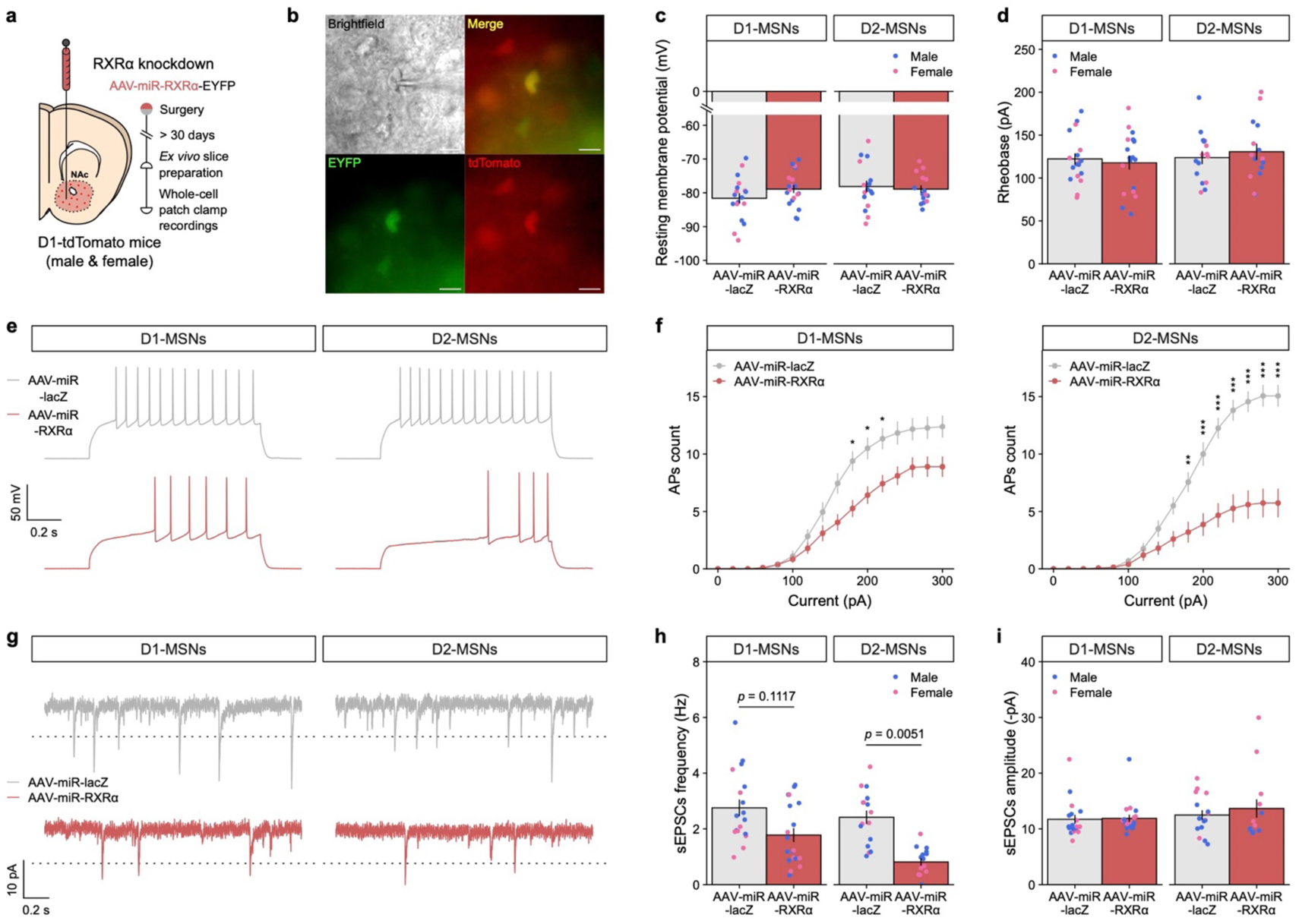
RXRα modulates intrinsic excitability of and spontaneous synaptic activity in both D1- and D2-MSNs. **a**, Experimental design of cell-type-specific RXRα knockdown *ex vivo* electrophysiology study. Male and female Drd1-tdTomato mice (n = 4-5/group) were injected in NAc with an AAV construct expressing a knockdown miRNA-sequence targeted against *Rxra* or a control *Lacz* sequence, then euthanized at least 30 days later. Acute *ex vivo* NAc slices were prepared before whole-cell patch-clamp recordings of tdTomato^+^ (D1-MSNs) or tdTomato^−^ (D2-MSNs) cells (n = 68 neurons total). **b**, Representative picture of a patched AAV-infected D1-tdTomato^+^ neuron. Scale bar is 20 µm. **c**, Resting membrane potential was unaffected by RXRα knockdown in either D1- or D2-MSNs (LMM-ANOVA: main effect of Virus: F_1,4.978_ = 0.8204, *p* = 0.4068; interaction Virus:CellType: F_1,57.479_ = 1.5532, *p* = 0.2177, followed by Sidak’s *post hoc* tests). **d**, Rheobase, the minimal injected current required to trigger an action potential, was also unaffected by RXRα knockdown in either D1- or D2-MSNs (LMM-ANOVA: main effect of Virus: F_1,5.79_ = 0.0207, *p* = 0.8904; interaction Virus:CellType: F_1,55.6_ = 2.0595, *p* = 0.1569, followed by Sidak’s *post hoc* tests). **e**, Representative membrane responses from D1- and D2-MSNs in response to a 200 pA current injection, with or without RXRα knockdown. **f**, The number of evoked action potentials (AP) in response to increasing depolarizing current steps was blunted by RXRα knockdown in both D1- (left) and D2-MSNs (right), suggesting reduced excitability (LMM-ANOVA: interaction Virus:Current: F_1,1012_ = 293.59, *p* < 2e^−16^; interaction Virus:Current:CellType: F_1,1012_ = 61.4425, *p* = 1.15e^−14^, followed by Sidak’s *post hoc* tests: * *p* < 0.05, ** *p* < 0.01, *** *p* < 0.001). **g**, Representative voltage-clamp recordings of NAc D1- or D2-MSNs highlighting spontaneous Excitatory Post-Synaptic Currents (sEPSCs), with or without RXRα knockdown. Dotted line represents the 8 pA deviation from baseline threshold for sEPSC detection. **h**, RXRα knockdown reduced the frequency of sEPSCs in both D1- and D2-MSNs (LMM-ANOVA: main effect of Virus: F_1,5.22_ = 16.8442, *p* = 0.008509; interaction Virus:CellType: F_1,57.144_ = 1.619, *p* = 0.198575, followed by Sidak’s *post hoc* tests). **i**, RXRα knockdown did not alter the amplitude of sEPSCs in either D1- or D2-MSNs (LMM-ANOVA: main effect of Virus: F_1,5.26_ = 0.5445, *p* = 0.4922; interaction Virus:CellType: F_1,56.191_ = 0.3214, *p* = 0.5730, followed by Sidak’s *post hoc* tests). Bar graphs and line graphs represent mean ± sem after combining male and female data.

### RXRα cell-autonomously adjusts neuronal excitability in D1- and D2-MSNs

The next step was thus to investigate the cell-type-specific physiological consequences of RXRα-dependent transcriptional control of excitability- and plasticity-related genes. We switched to an RXRα loss-of-function knockdown strategy, and developed miRNA constructs targeted against the *Rxra* coding sequence or a control *Lacz* sequence (not expressed in mammals), both of which were fused to an EYFP reporter, and packaged into AAVs (AAV-miR-RXRα-EYFP or AAV-miR-lacZ-EYFP). RXRα knockdown efficiency in virally-infected tissue punches was at least 50% at both the RNA and protein levels (Extended Data Fig. S2). These constructs were then infused into the NAc of D1-tdTomato male and female mice, which express a tdTomato fluorophore under the promoter of the *Drd1* gene (Fig. 4a). This strategy allowed us to record from AAV-infected (EYFP^+^) D1-MSNs (tdTomato^+^) or D2-MSNs (tdTomato^−^) (Fig. 4b). In both cell types, reducing RXRα levels did not affect resting membrane potential (Fig. 3c, LMM-ANOVA: main effect of Virus: F_1,4.978_ = 0.8204, *p* = 0.4068; interaction Virus:CellType: F_1,57.479_ = 1.5532, *p* = 0.2177) nor rheobase (Fig. 3d, LMM-ANOVA: main effect of Virus: F_1,5.79_ = 0.0207, *p* = 0.8904; interaction Virus:CellType: F_1,55.6_ = 2.0595, *p* = 0.1569). Of note, average values for those two metrics were in line with previous reports ^39^. When measuring the number of action potentials evoked in patched MSNs upon increasing steps of current injection, we found strikingly dampened responses at higher currents in both cell types with RXRα knockdown (Fig. 4e,f, LMM-ANOVA: interaction Virus:Current: F_1,1012_ = 293.59, *p* < 2e^−16^), indicating blunted intrinsic excitability. At the synaptic level, frequency (Fig. 4g,h, LMM-ANOVA: main effect of Virus: F_1,5.22_ = 16.8442, *p* = 0.008509; interaction Virus:CellType: F_1,57.144_ = 1.619, *p* = 0.198575) but not amplitude (Fig. 4g,i, LMM-ANOVA: main effect of Virus: F_1,5.26_ = 0.5445, *p* = 0.4922; interaction Virus:CellType: F_1,56.191_ = 0.3214, *p* = 0.5730) of spontaneous excitatory post-synaptic currents (sEPSCs) was reduced in both cell types. RXRα‘s influence on intrinsic excitability was stronger in D2-MSNs (LMM-ANOVA: interaction Virus:Current:CellType: F_1,1012_ = 61.4425, *p* = 1.15e^−14^), and trended to be stronger for sEPSCs frequency as well (Sidak’s *post hoc* tests: D1-MSNs, *p* = 0.1117; D2-MSNs, *p* = 0.0051, although interaction Virus:CellType was not significant). These electrophysiological results therefore substantiate that the transcriptional programs controlled by RXRα indeed have the capacity to modulate intrinsic excitability and synaptic function of both MSN subtypes – likely via changes in the channel and receptor repertoires expressed at the postsynaptic membrane – and that RXRα could thus be critical in calibrating neuronal sensitivity to drug stimuli.

### Manipulating RXRα levels in NAc influences sensitivity to drug reward contextual learning

Our final efforts were thence to characterize the behavioral consequences of RXRα-mediated transcriptional regulation of MSN excitability. To assess whether NAc RXRα levels alter the ability of an animal to associate predictive contextual cues with the rewarding properties of drugs of abuse, we used an unbiased conditioned place preference (CPP) assay in male and female mice. HSV-mediated RXRα overexpression in all NAc neuronal types increased CPP formation for a subthreshold (5 mg/kg) dose of cocaine, which did not induce a preference in control animals (Fig. 5a, LMM-ANOVA: interaction Test:Virus: F_1,44_ = 10.1557, *p* = 0.002647, followed by Sidak’s *post hoc* tests). This effect generalized to other drug reinforcers, as it was also observed for an intermediate (7.5 mg/kg) dose of morphine (Fig. 5b, LMM-ANOVA: interaction Test:Virus: F_1,43_ = 9.4374, *p* = 0.003681, followed by Sidak’s *post hoc* tests). The impact of RXRα on cocaine reward was bidirectional, as AAV-mediated RXRα knockdown produced the opposite effect – a decrease in CPP for a higher (10 mg/kg) dose of cocaine (Fig. 5c, LMM-ANOVA: interaction Test:Virus: F_1,41_ = 5.1204, *p* = 0.02901, followed by Sidak’s *post hoc* tests). Because D1- and D2-MSNs have been shown to play opposite roles in drug-reward contextual learning ^33^, including in this specific CPP task ^31, 32^, we hypothesized that RXRα-mediated modulation of their intrinsic excitability would exert opposite effects on cocaine CPP. We found that D1-MSN-specifc (Fig. 5d, LMM-ANOVA: interaction Test:Virus: F_1,44_ = 28.4447, *p* < 0.0001, followed by Sidak’s *post hoc* tests) but not D2-MSN-specific (Fig. 5e, LMM-ANOVA: interaction Test:Virus: F_1,44_ = 0.0813, *p* = 0.7768) RXRα overexpression increased subthreshold cocaine CPP, indicating that net bulk tissue effects (Fig. 5a) are likely triggered through RXRα action in D1-MSNs. These experiments corroborate the correlative data shown in Fig. 1, as they causally establish that differential NAc RXRα levels control sensitivity to drug reward, with higher and lower RXRα levels respectively promoting and lessening drug reward learning.

**Fig. 5.**
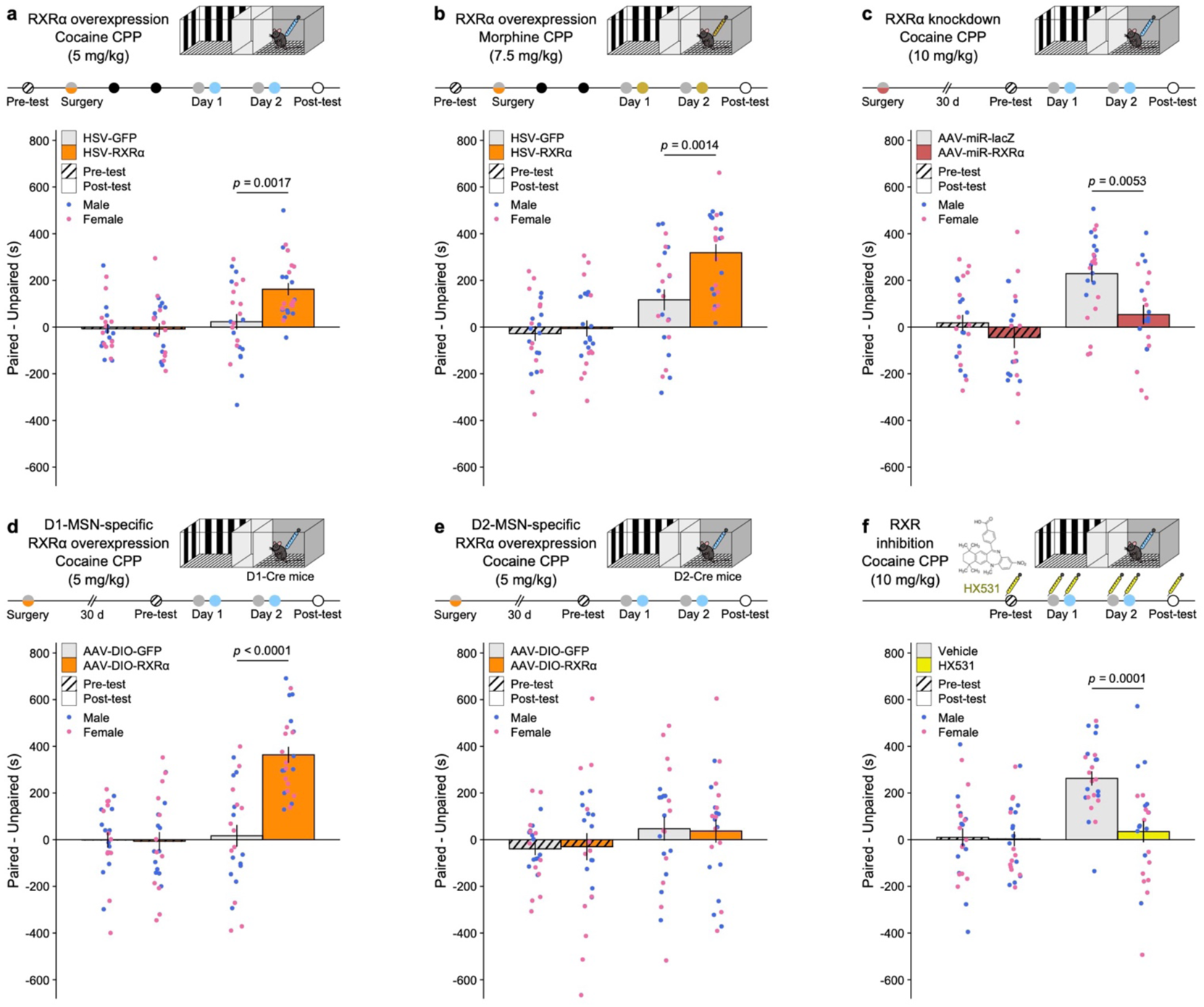
RXRα bidirectionally and cell-type-specifically regulates dose sensitivity in drug-reward associative learning. Both male and female mice were used in all experiments (n = 11-12/group). **a**, HSV-mediated RXRα overexpression in NAc increased conditioned place preference (CPP) for a subthreshold (5 mg/kg) dose of cocaine (LMM-ANOVA: interaction Test:Virus: F_1,44_ = 10.1557, *p* = 0.002647, followed by Sidak’s *post hoc* tests). **b**, HSV-mediated RXRα overexpression in NAc increased CPP for a 7.5 mg/kg dose of morphine (LMM-ANOVA: interaction Test:Virus: F_1,43_ = 9.4374, *p* = 0.003681, followed by Sidak’s *post hoc* tests). **c**, AAV-mediated RXRα knockdown in NAc decreased CPP for a high (10 mg/kg) dose of cocaine (LMM-ANOVA: interaction Test:Virus: F_1,41_ = 5.1204, *p* = 0.02901, followed by Sidak’s *post hoc* tests). **d**, AAV-mediated NAc D1-MSN-specific RXRα overexpression increased CPP for a subthreshold (5 mg/kg) dose of cocaine (LMM-ANOVA: interaction Test:Virus: F_1,44_ = 28.4447, *p* < 0.0001, followed by Sidak’s *post hoc* tests). **e**, AAV-mediated NAc D2-MSN-specific RXRα overexpression did not increase CPP for a subthreshold (5 mg/kg) dose of cocaine (LMM-ANOVA: interaction Test:Virus: F_1,44_ = 0.0813, *p* = 0.7768, followed by Sidak’s *post hoc* tests). **f**, Pharmacological inhibition of RXRα via systemic administration of RXR antagonist HX531 (10 mg/kg) decreased CPP for a high (10 mg/kg) dose of cocaine (LMM-ANOVA: interaction Test:Virus: F_1,44_ = 13.4504, *p* = 0.0006572, followed by Sidak’s *post hoc* tests). Bar graphs represent mean ± sem after combining male and female data.

### Systemic RXRα pharmacological inhibition weakens cocaine reward

Furthermore, intraperitoneal injection of the RXR-family antagonist HX531 ^40^, at a dose (20 mg/kg) known to interfere with striatal processes like amphetamine-induced hyperlocomotion ^41^ and haloperidol-induced dyskinesias ^42^, blocked cocaine CPP (Fig. 5f, LMM-ANOVA: interaction Test:Virus: F_1,44_ = 13.4504, *p* = 0.0006572, followed by Sidak’s *post hoc* tests). This experiment supports the possibility that systemically targeting RXRα is a therapeutically accessible approach to combat drug addiction.

### RXRα enhances cocaine reinforcing efficacy in goal-directed tasks

CPP measures of associative learning have been interpreted as a direct function of a stimulus’s rewarding value, however, they also incorporate essential elements of attention, saliency and learning mechanisms ^43^. To better dissociate changes in dose sensitivity and reinforcing efficacy from these other factors, especially from learning components, we combined HSV-mediated RXRα overexpression with a self-administration behavioral economics threshold procedure (Fig. 6a) that assesses an animal’s motivation to self-administer a reinforcer in the face of increasing cost by generating a dose-response curve within subject within a single session ^44–46^. This procedure was utilized in male rats – where it has been best validated ^47^ – to also further confirm the role of RXRα in another model species. After training on a FR1 schedule and HSV infusion, we first observed that HSV-RXRα rats reduced their total drug intake in post-surgery FR1 sessions (Fig. 6b, unpaired Welch’s *t*-test: *t*_8.5723_ = 2.3018, *p* = 0.04824). However, during behavioral economics testing, they exhibited significantly higher responding for low (1-5 µg) doses of cocaine (Fig. 6c, LMM-ANOVA: interaction Dose:Virus: F_9,126_ = 5.4513, *p* = 0.00000261, followed by Sidak’s *post hoc* tests), indicating an upward and leftward shift of the dose-response function after RXRα overexpression. Next, data were plotted as a demand curve (Fig. 6d), with individual consumption levels as a function of price (number of active lever presses required for 1 mg of cocaine). Basic economic principles were then applied through mathematical modeling of individual demand curves (Fig. 6e) to extrapolate independent measures of consumption, motivation and demand elasticity ^46, 48^. Consistent with FR1 consumption data (Fig. 6b), both consumption at low effort Q_0_ (Fig. 6f, unpaired Welch’s *t*-test: *t*_13.988_ = 1.9504, *p* = 0.07145) and at maximum effort Q_max_ (Fig. 6g, unpaired Welch’s *t*-test: *t*_13.985_ = 1.8907, *p* = 0.07956) showed trends toward a decrease with RXRα overexpression. By contrast, and consistent with dose-response data (Fig. 6c), motivation metrics such as the price of maximal responding after which consumption decreases P_max_ (Fig. 6h, unpaired Welch’s *t*-test: *t*_8.6384_ = −3.2059, *p* = 0.01131) and the maximal behavioral output at that price O_max_ (Fig. 6i, unpaired Welch’s *t*-test: *t*_8.0425_ = −2.3817, *p* = 0.04427) were both significantly increased by RXRα overexpression. Finally, RXRα overexpression in NAc reduced demand elasticity α, which indicates how sensitive demand is to price and can be interpreted as an inverse indicator of the essential value of the reinforcer used; our findings show lower α suggesting higher cocaine essential value after RXRα overexpression (Fig. 6j, unpaired Welch’s *t*-test: *t*_12.216_ = 2.4123, *p* = 0.03245). It is important to emphasize that all these metrics were equivalent across animals when tested before HSV surgery and group allocation (Extended Data Fig. S5).

**Fig. 6.**
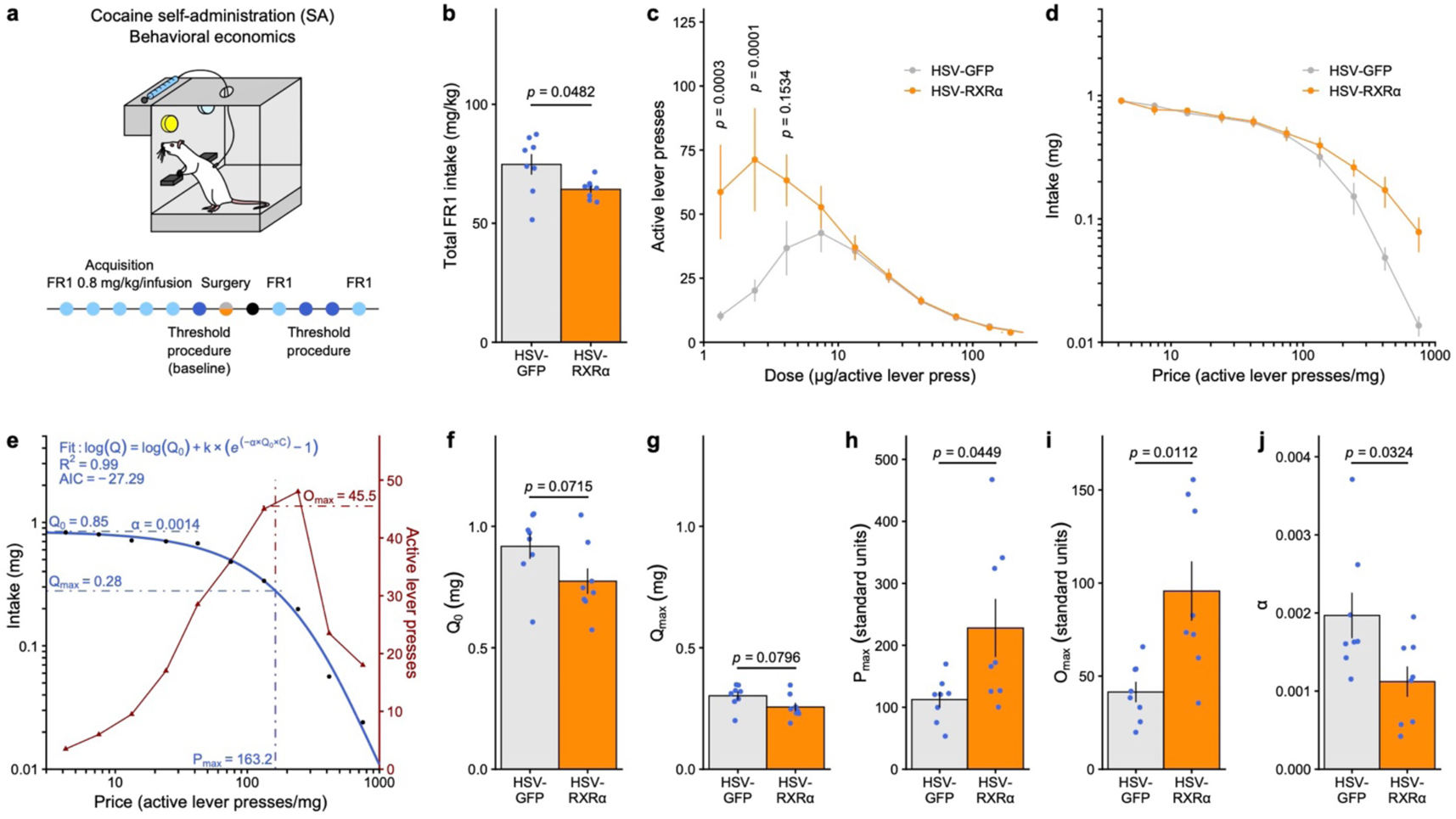
RXRα overexpression in NAc increases motivation to self-administer low cocaine doses. **a**, Experimental design for behavioral economics testing. Male rats (n = 8/group) were trained to self-administer cocaine (0.8 mg/kg/infusion) on an FR1 schedule of reinforcement before being injected in NAc with an HSV overexpressing RXRα or a control GFP sequence. Animals then went through two behavioral tasks: the threshold procedure to assess motivation and FR1 to assess consumption. **b**, Total cocaine intake in FR1 sessions was significantly reduced by RXRα overexpression (unpaired Welch’s *t*-test: *t*_8.5723_ = 2.3018, *p* = 0.04824). **c**, Dose response curves in the threshold procedure showed increased responding for low doses of cocaine after RXRα overexpression (LMM-ANOVA: interaction Dose:Virus: F_9,126_ = 5.4513, *p* = 0.00000261, followed by Sidak’s *post hoc* tests). **d**, Averaged demand curves showed a similar trend of increased cocaine intake at higher prices (LMM-ANOVA: interaction Price:Virus: F_9,126_ = 1.2085, *p* = 0.2955, followed by Sidak’s *post hoc* tests). **e**, Representative example of task performance in the threshold procedure (from a HSV-GFP control), highlighting mathematical demand curve fitting and extraction of behavioral economics parameters. Goodness of fit was assess by computing a pseudo-R^2^ coefficient and the Akaike Information Criterion (AIC) for each individual fit. **f**, Consumption at low effort Q_0_ (unpaired Welch’s *t*-test: *t*_13.988_ = 1.9504, *p* = 0.07145) and **g**, consumption at maximum effort Q_max_ (unpaired Welch’s *t*-test: *t*_13.985_ = 1.8907, *p* = 0.07956) both trended toward a decrease after RXRα overexpression. Motivation metrics **h**, P_max_ (unpaired Welch’s *t*-test: *t*_8.6384_ = −3.2059, *p* = 0.01131) and **i**, O_max_ (unpaired Welch’s *t*-test: *t*_8.0425_ = −2.3817, *p* = 0.04427) were both increased by RXRα overexpression. **j**, Demand elasticity α was reduced by RXRα overexpression (unpaired Welch’s *t*-test: *t*_12.216_ = 2.4123, *p* = 0.03245). Bar graphs and line graphs represent mean ± sem.

To summarize, RXRα overexpression in NAc reduced cocaine consumption but increased sensitivity to and motivation for low cocaine doses, as well as cocaine value. Decorrelation between consumption and motivation is precedented ^44, 49, 50^, and here we propose that rats with increased RXRα NAc levels consume less cocaine precisely because they are in fact more sensitive to its reinforcing properties and rewarding value, and thus consume less drug to maintain a given hedonic set point. Extending upon our CPP data (Fig. 5), these operant behavioral findings demonstrate that RXRα dictates individual sensitivity to drug rewards through direct modulation of their reinforcing efficacy.

## DISCUSSION

In this study, we identify RXRα as a novel key mediator of the NAc transcriptional mechanisms underlying sensitivity to drug reinforcers. Using a combination of viral approaches, we causally demonstrate a positive association between individual RXRα levels in NAc and addiction-relevant molecular, cellular and behavioral features: the higher levels at which select NAc cell types express RXRα, the more excitable they are (Fig. 4) and the more responsive to low drug doses the animal is (Fig. 5, 6). Our data also support the idea that these effects are transcriptionally mediated at baseline before drug exposure (Fig. 2, 3), and are also recruited as important mediators of the transcriptional plasticity mechanisms induced after acute or repeated drug exposure (Fig. 1, 2).

Transcriptional processes represent a fundamental building block of neuronal function and plasticity, including after drug exposure ^4, 10^. In that context. RXRα is particularly well-suited to fine-tune these transcriptional programs in a very specific yet stimulus-dependent manner, because of its ability to function either as homodimers or as an obligatory heterodimerization partner for several other nuclear receptors, like brain-expressed retinoic acid receptors (RARs), liver X receptors (LXRs), peroxisome proliferator-activated receptors (PPARs), vitamin D receptors (VDRs) and thyroid hormone receptors (TRs) among many others ^13, 34, 35, 51^. In addition to bridging together different signaling pathways, such heterodimers can be either permissive or repressive for transcription of target genes depending on subsequent recruitment of distinct co-regulators and epigenetic enzymes ^13, 34, 35^. This posits RXRα as a potential integrative signaling hub at key gene promoters to subtly dictate locus-specific gene expression upon activation of other, well-established, generic drug-evoked signaling cascades and transcription factors such as CREB ^12^. Consistent with this idea, we have shown that RXRα motifs exist in close proximity to CREB binding sites at promoters of select cocaine-regulated genes ^11^. Other promising candidate RXRα binding partners include 1) RXRγ, which is known to contribute to striatal- and dopamine-dependent locomotion ^18^ and depressive-like ^19–21^ behaviors, 2) NURR77/NR4A1, an IEG and orphan receptor associated with cocaine ^52^ and stress ^53^ behaviors that we here found positively correlated with RXRα expression (Fig. 2e), 3) RARs and PPARs because the enzymes that deliver retinoic acid to RAR/RXR or PPAR/RXR complexes – CRABP2 and FABP5 respectively – differentially contribute to addiction- ^15, 16^ and depression-related ^17^ behaviors, and 4) VDRs, as vitamin D deficiency was recently linked with opioid addiction ^54^. Ultimately, a finer dissection of RXRα binding partners could better explain some of the cell-type-, region-, drug- and drug-regimen-specificity of drug-induced transcriptional programs, as these likely result not from the recruitment of one given transcription factor but rather of several factors that could be partly harmonized by RXRα, among others. Future research is also needed to understand how baseline RXRα-dependent excitability-related transcriptional programs contribute to drug-induced transcriptional activation and plasticity by elucidating potential coordinated shifts in RXRα binding partners and their target binding sites upon drug exposure.

By definition, activation of specific nuclear receptor complexes is dependent on proper recognition of their respective ligands ^13, 34, 35, 51^, making ligand dynamics a critical component for transducing environmental stimuli into transcriptional responses. However, the exact nature of the endogenous brain ligand(s) for RXR family members including RXRα – termed rexinoids – remains debated, partially because of technical difficulties in detecting and studying naturally-occurring retinoids and rexinoids ^34, 51^. *All-trans*-retinoic acid (atRA), the undisputed physiological ligand for RARs ^55^, does not bind RXRs ^55^. *9-cis*-retinoic acid (9CRA), which binds RXRs with nanomolar affinity *in vitro* ^55^, was considered as an RXR endogenous ligand until several reports failed to detect endogenous 9CRA levels *in vivo* ^51, 56^. Several other molecules have since been proposed, but to date the two strongest candidates for rexinoid activity in brain are 1) free fatty acids metabolites of docosahexaenoic acid (DHA), especially unesterified DHA, and 2) dihydroretinoid metabolites, especially *9-cis-13,14*-dihydroretinoic acid (9CDHRA). On the one hand, DHA is abundant in the mammalian brain ^57^ (although whether in forms and at concentrations sufficient to induce transactivation is questioned ^51^), has been shown to promote cortical excitability and synaptic architecture in a RXRα-dependent manner ^29^, and DHA supplementation can reverse depressive-like behavior in an RXRγ-dependent manner ^20^. On the other hand, 9CDRHA is detected in brain tissue at high levels sufficient for RXR binding ^56^, and 9CDHRA supplementation also reversed depressive-like behavior in a RXRγ-dependent manner ^21^. Whether DHA, 9CDRHA or other related molecules binds RXRα in NAc, as well as the exact kinetics of their action (baseline levels, but also hypothetical release temporal dynamics), and whether these are affected by drug exposure, remain to be elucidated, and would greatly benefit from improved resolution of metabolomics techniques.

One shared property of rexinoids is that they or their precursors can be nutritionally provided ^19–21, 51, 57^. This has great preclinical relevance for several other pathologies linked to RXR signaling including Alzheimer’s disease, Parkinson’s disease, multiple sclerosis and diabetes ^34, 35, 51^ and has sparked interest in testing phyto-pharmacologically derived rexinoids or *de novo* synthetic RXR ligands to clinically combat or prevent these conditions ^34, 35, 51^. Here we show that inhibition of RXRs using a systemically administered RXR antagonist (HX531) reduced cocaine-induced associative learning (Fig. 5f), consistent with previous reports showing that HX531 blocked amphetamine-induced hyperlocomotion in a NURR77/NR4A1-dependent manner ^41^ – another hint at further studies of RXR/NURR77 dimers in psychostimulant action (see above). Moreover, this inhibitor has been shown to block RXR/RXR, RAR/RXR and PPARγ/RXR, but not PPARα/RXR heterodimers ^40^, highlighting rexinoid drugs as a potential platform from which to derive pharmacological compounds with increased specificity for certain RXR-containing nuclear receptor dimers – *i.e.*, for some target pathways, cell type or condition. Finally, because full cortical and hippocampal RXRα knockout mice behaved normally on an extensive battery of baseline behavioral tests ^29^, our preliminary data illustrates that systemically or nutritionally targeting NAc RXRα using rexinoids might be leveraged to design improved pharmacological treatments for drug addiction with tolerable side effects.

In conclusion, this work represents a meaningful advance in the study of drug reinforcement mechanisms and their contribution to drug addiction. We single out a novel and putatively druggable molecule, RXRα, that transcriptionally regulates larger gene networks to fine-tune the excitability of NAc cell types and in turn individual sensitivity to drugs of abuse. This work also underscores the effectiveness of bioinformatically harnessing large-scale datasets ^11^ as part of the effort to unravel innovative answers to highly complex brain maladaptations from both fundamental and translational standpoints.

## METHODS

### Animals

Animals were housed in the animal facilities at Mount Sinai. Male and female C57BL/6J mice (8-16 weeks old, 20-30 g, The Jackson Laboratory) were maintained on a 12:12 h light/dark cycle (07:00 lights on; 19:00 lights off) and were provided with food and water *ad libitum*. Male Sprague-Dawley rats (300-400g) were maintained on a 12:12 h reverse light/dark cycle (07:00 lights off; 19:00 lights on) and restricted to 95% of free-feeding body weight to improve consistency of the self-administration behavior. Transgenic mouse lines (D1-Cre: MGI:3836633, D2-Cre: MGI:3836635, D1-tdTomato: MGI:4360387, ^fl/fl^eGFP::L10a: IMSR_JAX:022367) were bred in-house on a C57BL/6J background. All animals were maintained according to the National Institutes of Health guidelines for Association for Assessment and Accreditation of Laboratory Animal Care accredited facilities. All experimental protocols were approved by the Institutional Animal Care and Use Committee at Mount Sinai.

### Drug treatments

Cocaine HCl (from the National Institute on Drug Abuse) was diluted in 0.9% NaCl saline solution (ICU Medical) and injected intraperitoneally at 20, 10 or 5 mg/kg. Morphine SO_4_ (from the National Institute on Drug Abuse) was diluted in 0.9% NaCl saline solution (ICU Medical) and injected subcutaneously at 7.5 mg/kg. HX531 (Tocris, Cat. No. 3912) was first diluted in DMSO (Sigma), then in Dulbecco’s Phosphate Buffered Saline (PBS, Gibco) to a 1% DMSO final concentration and injected intraperitoneally at 20 mg/kg.

### Viral reagents

For HSV-mediated RXRα overexpression, an RXRα insert consisting of a codon-optimized mouse sequence reverse-translated from RXRα mouse protein sequence was synthesized *de novo* by Genewiz into a pENTR plasmid, then sub-cloned downstream of a IE4/5 promoter into a bicistronic p1005 plasmid, which also expresses GFP under a separate CMV promoter, using Gateway LR Clonase Kit (Invitrogen). Packaging into HSVs of the overexpression (HSV-RXRα-GFP) and empty control (HSV-GFP) vectors was performed at the Gene Delivery Technology Core of Massachusetts General Hospital in Boston, Massachusetts, USA. For Cre- dependent AAV-mediated RXRα overexpression, the RXRα insert was excised from the p1005 plasmid and sub-cloned into an AAV2 expression backbone downstream of a CAG promoter and of an EGFP reporter sequence separated by a P2A cleavage sequence. The EGFP-P2A-RXRα sequence was flanked by loxP site to only express the transgene upon Cre-dependent recombination. Cloning and then packaging into AAV9 capsids of the overexpression (AAV9-CAG-DIO-EGFP-P2A-RXRα) and control (AAV9-CAG-DIO-EGFP) vectors were performed at the Duke University Viral Vector Core in Durham, North Carolina, USA. For AAV-mediated RXRα knockdown, miRNA plasmids were prepared using the BLOCK-iT Pol II miR RNAi Expression Vector Kit (ThermoFisher Scientific) with top and bottom sequences targeting mouse *Rxra* cDNA (Top: 5’ - TGC TG<u>A GAT GTT GGT AAC AGG GTC AT</u>G TTT TGG CCA CTG ACT GAC <u>ATG ACC CTT ACC AAC</u> <u>ATC T</u> - 3’; Bottom: 5’ – CCT GAG ATG TTG GTA AGG GTC ATG TCA GTC AGT GGC CAA AAC C - 3’) or a control *Lacz* sequence (Top: 5’ – TGC TG<u>A AAT CGC TGA TTT GTG TAG TC</u>G TTT TGG CCA CTG ACT GAC <u>GAC TAC ACA TCA GCG ATT T</u> - 3’; Bottom: 5’ – CCT GAA ATC GCT GAT GTG TAG TCG TCA GTC AGT GGC CAA AAC GAC TAC ACA AAT CAG CGA TTT C - 3’) obtained from Invitrogen. The miRNA plasmids were then sub-cloned into an AAV2 expression backbone downstream of a CMV promoter and of an EYFP reporter sequence. Cloning and then packaging into AAV9 capsids of the knockdown (AAV9-CMV-EYFP-miR-RXRα) and control (AAV9-CMV-EYFP-miR-lacZ) vectors were performed by the Duke University Viral Vector Core in Durham, North Carolina, USA. All viral vectors were validated *in vivo* (Extended Data Fig. S2).

### Stereotaxic surgeries

Mice or rats were anesthetized with an intraperitoneal bolus of ketamine (100 mg/kg) and xylazine (10 mg/kg), then head-fixed in a stereotaxic apparatus (Kopf Instruments). Syringe needles (33G, Hamilton) were used to bilaterally infuse 1 µl of virus at a 0.1 µl/min flow rate. Needles were kept in place for 10 minutes after injection before being retracted to allow for virus diffusion. Viruses were used at the following titers: HSV-GFP and HSV-RXRα-GFP at ± 1 × 10^8^ IU/mL, AAV9-CAG-DIO-EGFP-P2A-RXRα and AAV9-CAG-DIO-EGFP at ± 1 × 10^12^ VG/mL, AAV9-CMV-EYFP-miR-RXRα and AAV9-CMV-EYFP-miR-lacZ at ± 5 × 10^12^ VG/mL. Coordinates for NAc were as follows, from Bregma: AP + 1.6 mm, ML + 1.6 mm, DV – 4.5 mm, 10° angle for mice, and AP + 1.9 mm, ML + 2.8 mm, DV – 7.2 mm, 10° angle for rats.

### RNA extraction and quantitative real-time PCR

Mouse brains were collected after cervical dislocation and followed by rapid bilateral NAc punch dissections from 1 mm-thick coronal brain sections using a 14G needle and frozen on dry ice. RNA extraction was performed using the RNeasy Micro Kit (Qiagen) following manufacturer instructions. RNA 260/280 ratios of 2 were confirmed using spectroscopy, and reverse transcription was achieved using the iScript cDNA Synthesis Kit (BioRad). Quantitative PCR using PowerUp SYBR Green (Applied Biosystems) was used to quantify cDNA using an Applied Biosystems QuantStudio 5 system. Each reaction was performed in triplicate and relative expression was calculated relative to the geometric average of 3 control genes (*Ppia*, *Tbp*, *Rpl38*) according to published methods ^58^. Sequences of all used primers are available in Extended Data Table S4.

### Protein extraction and Western blotting

Mouse brains were collected after cervical dislocation and followed by rapid bilateral NAc punch dissections from 1 mm-thick coronal brain sections using a 14G needle and frozen on dry ice. For whole tissue extracts, frozen NAc samples were homogenized and then incubated for 30 min with agitation in 200 µL of ice-cold RIPA buffer (10 mM Trizma Base, 150 mM NaCl, 1 mM EDTA, 0.1% SDS, 1% Triton-X-100, 1% sodium deoxycholate, pH 7.4, complemented with protease and serine/threonine and tyrosine phosphatase inhibitors), before 5 cycles of 20 s on/off sonication using a Bioruptor (Diagenode). Samples were centrifuged for 15 min at 14,000 g to pellet insoluble debris and lipids, and supernatant was transferred to new tubes. For nuclear fractions, frozen NAc samples were first subjected to subcellular fractionation using our established procedures ^44^. Briefly, frozen NAc samples were homogenized in 150 µL of HEPES-buffered sucrose (0.32 M sucrose, 4 mM HEPES, pH 7.4, complemented with protease and serine/threonine and tyrosine phosphatase inhibitors) before a 10 min centrifugation at 1000 g at 4°C to pellet nuclei, followed by 3 washes in HEPES-buffered sucrose and finally by RIPA extraction (see above). For both whole tissue and nuclear extracts, protein concentration was quantified using a Pierce BCA Protein Assay Kit (ThermoFisher Scientific). SDS-PAGE protein separation and Western blotting were performed according to manufacturer instructions and our published protocols ^44^. Equal amounts of proteins were mixed !3-mercaptoethanol-supplemented Laemmli buffer (BioRad), heated to 95°C for 5 min before being separated by SDS-PAGE with Criterion Precast Gels (4 –15% Tris-HCl; BioRad) and transferred onto Immun-Blot PVDF 0.2 µm (BioRad) membranes. Membranes were blocked in Tris-buffered saline containing 5% bovine serum albumin (Sigma) and 0.1% Tween-20 (Fisher Bioreagents) for 1 h at room temperature. Primary antibodies (full list and concentrations used available in Extended Data Table S5) were diluted in blocking solution and incubated overnight at 4°C. After washing, membranes were incubated with anti-mouse (#PI-2000, Vector Laboratories) or antirabbit (#PI-1000, Vector Laboratories) peroxidase-conjugated secondary antibodies diluted 1:50,000 in blocking solution for 2 h, washed thoroughly, and developed using SuperSignal West Dura Substrate (ThermoFisher Scientific). Quantification was performed by densitometry using ImageJ (U.S. National Institutes of Health). Protein levels were normalized to actin for whole tissue extracts or to actin and TBP for nuclear extracts. Between primary antibodies, membranes were stripped using Restore Plus Stripping Buffer (ThermoFisher Scientific). Male and female samples were run on separate gels and later normalized to their respective controls before being pooled for analysis. Unmodified Western blots scans are provided as Source Data.

### Nuclei purification and Fluorescence-Activated Nuclei Sorting (FANS)

Mouse brains were collected after cervical dislocation and followed by rapid bilateral NAc punch dissections of virally-infected tissue under fluorescent light from 1 mm-thick coronal brain sections using a 14G needle and frozen on dry ice. To obtain a single nuclei suspension, frozen NAc samples were homogenized in 4 mL of low-sucrose lysis buffer (0.32 M sucrose, 5 mM CaCl_2_, 3 mM Mg(Ace)_2_, 0.1 mM EDTA, 10 mM Tris-HCl) using a large clearance then a small clearance pestle of a glass dounce tissue grinder (Kimble Kontes). Homogenates were filtered through a 40 µm cell strainer (Pluriselect) into ultracentrifuge tubes (Beckman Coulter), underlaid with 5 mL of high-sucrose solution (1.8 M sucrose, 3 mM Mg(Ace)_2_, 1 mM DTT, 10 mM Tris-HCl) and centrifugated at 24,000 rpm for 1 h at 4°C in a SW41Ti Swinging-Bucket Rotor (Beckman Coulter). Supernatant was discarded and nuclei pellets were re-suspended in 800 µL of PBS. DAPI was added at a 1:5000 dilution. Nuclei were sorted on a BD FACS Aria II three-laser device with a 70 µm nozzle. Gating strategy from representative sorts are visualized in Extended Data Fig. S4. Briefly, debris and doublets were excluded using FSC and SSC filters, nuclei were then selected as DAPI-positive (Violet1-A laser) events, and finally GFP-positive nuclei (Blue1-A laser) were sorted directly into TRIzol LS (Ambion) and flash frozen. Between 30,000 and 60,000 nuclei were recovered for each sample.

### RNA-sequencing (RNAseq)

For bulk tissue RNAseq, mouse brains were collected after cervical dislocation and followed by rapid bilateral NAc punch dissections of virally-infected tissue under fluorescent light from 1 mm-thick coronal brain sections using a 14G needle and frozen on dry ice. RNA extraction was performed using the RNeasy Micro Kit (Qiagen) following manufacturer instructions. Sequencing libraries were prepared by Genewiz/Azenta with polyA selection and sequenced with Genewiz/Azenta on an Illumina HiSeq 4000 machine using a 2 × 150 bp pair-end read configuration to a minimum depth of 20 million reads per sample. For cell-type-specific RNAseq, RNA was extracted from frozen TRIzol LS (Ambion) homogenates using the Direct-zol RNA Microprep Kit (Zymo Research) following manufacturer instructions. Ribo-depleted sequencing libraries were prepared with the SMARTer Stranded Total RNA-Seq Kit v2 - Pico Input Mammalian (TaKaRa Biotech) following manufacturer instructions and sequenced with Genewiz/Azenta on an Illumina NovaSeq S4 machine using a 2 × 150 bp pair-end read configuration to a minimum depth of 40 million reads per sample. For both experiments, quality control was performed using FastQC (www.bioinformatics.babraham.ac.uk/projects/fastqc/), adapter contaminants were trimmed using Trimmomatic (github.com/usadellab/Trimmomatic), reads were aligned to a custom-built reference genome (RXRα and GFP plasmid sequences were added to GENCODE GRCm38 genome) using hisat2 (daehwankimlab.github.io/hisat2/), duplicate reads were removed using Picard MarkDuplicate tool (broadinstitute.github.io/picard/), and finally count matrices were generated using the featureCounts function of the subread package (subread.sourceforge.net/featureCounts.html) and a comprehensive gene annotation file obtained from GENCODE (GRCm38 vM23). Reads mapping to the plasmid codon-optimized *Rxra* overexpression sequence were added to reads mapping to the endogenous *Rxra* sequence (ENSMUSG00000015846). The top 30% most expressed annotated genes/features (highest normalized read counts average across all samples) were kept for subsequent analysis in order to filter out poorly expressed genes. Differential expression was analyzed in R v 4.0.2 using the *DESeq2* package ^59^. Of note, for the cell-type-specific RNAseq both males and females were used, and biological sex was included as a factor in the model design (∼ Sex + Virus). To confirm cell-type specificity, only GFP control samples were selected and differential expression was run between cell types (∼ Sex + CellType model). Significance cut-offs were of at least 15% expression fold change (|log_2_(FoldChange)| > log_2_(1.15)) and nominal *p* < 0.05, except in the cases when genes were examined individually, where the Benjamini-Hochberg corrected *p_adj_* was used. Gene lists and corresponding statistics are available in Extended Data Table S2 (D1-MSNs) and S3 (D2-MSNs). RRHO plots were generated using the *RRHO2* package ^60^ (github.com/RRHO2/RRHO2). Analysis code is available upon request.

### RNA fluorescent in situ hybridization (FISH)

Mice were transcardially perfused with a fixative solution containing 4% paraformaldehyde (PFA). Brains were post-fixed for 24 h in 4% PFA at 4°C. Sections of 20 µm thickness were cut in the coronal plane with a vibratome (Leica) and stored at −20°C in a cryoprotectant solution containing 30% ethylene glycol (v/v), 30% glycerol (v/v) and 0.1 M phosphate buffer. NAc slices were mounted on charged Superfrost Plus microscope slides (Fisher Scientific) and processed for RNA FISH using RNAscope Multiplex Fluorescent Reagent Kit v2 (ACD Bio) according to manufacturer instructions using mouse probes for *Rxra* (Mm-Rxra, #463121), *Drd1a* (Mm-Drd1a-C2, #406491-C2) and Drd2 (Mm-Drd2-C3, #406501-C3) transcripts. Sections were counterstained with DAPI and mounted using ProLong Gold Antifade Mountant (ThermoFisher Scientific). Confocal images (6-10 per animal, 1024 × 1024 pixels, 16 bits pixel depth) were acquired on a SP8 inverted confocal microscope (Leica) using a 40X objective. A custom-made automated ImageJ (U.S. National Institutes of Health) pipeline was used to extract channel intensity in every nucleus identified on DAPI staining (Extended Data Fig. S3a). Individual *Rxra* puncta were detected using ComDet v 0.5.4 (github.com/ekatrukha/ComDet). Analysis code is available upon request.

### Ex vivo slice electrophysiology

After at least 4 weeks of recovery from AAV surgery, male and female D1-Tomato mice were anesthetized using isoflurane. Brains were rapidly extracted, and coronal sections (250 µm) were prepared using a Compresstome (Precisionary Instruments) in cold (0-4°C) sucrose-based artificial cerebrospinal fluid (SB-aCSF) containing 87 mM NaCl, 2.5 mM KCl, 1.25 mM NaH_2_PO_4_, 4 mM MgCl_2_, 23 mM NaHCO_3_, 75 mM Sucrose, 25 mM Glucose. After recovery for 60 min at 32°C in oxygenated (95% CO_2_ / 5% O_2_) aCSF containing 130 mM NaCl, 2.5 mM KCl, 1.2 mM NaH_2_PO_4_, 2.4 mM CaCl_2_, 1.2 mM MgCl_2_, 23 mM NaHCO_3_, 11 mM Glucose, slices were kept in the same medium at room temperature for the rest of the day and individually transferred to a recording chamber continuously perfused at 2-3 mL/min with oxygenated aCSF. Patch pipettes (4-6 MO) were pulled from thin wall borosilicate glass using a micropipette puller (Sutter Instruments) and filled with a K-Gluconate-based intra-pipette solution containing 116 mM KGlu, 20 mM HEPES, 0.5 mM EGTA, 6 mM KCl, 2 mM NaCl, 4 mM ATP, 0.3 mM GTP (pH 7.2). Cells were visualized using an upright microscope with an IR-DIC lens and illuminated with a white light source (Olympus for Scientifica), and fluorescence visualized through eGFP and mCherry bandpass filters upon LED illumination through the objective (p3000^ULTRA^, CoolLed). Excitability was measured in current-clamp mode by injecting incremental steps of current (0-300 pA, +20 pA at each step). For recording of spontaneous Excitatory Post-Synaptic Currents (sEPSCs), NAc MSN neurons were recorded in voltage-clamp mode at −70mV and detected with a 8 pA threshold. Whole-cell recordings were performed using a patch-clamp amplifier (Axoclamp 200B, Molecular Devices) connected to a Digidata 1550 LowNoise acquisition system (Molecular Devices). Signals were low pass filtered (Bessel, 2 kHz) and collected at 10 kHz using the data acquisition software pClamp 11 (Molecular Devices). Electrophysiological recordings were extracted using Clampfit (Molecular Devices). All groups were counterbalanced by days of recording and all recordings were performed blind to experimental condition.

### Conditioned place preference (CPP)

Unbiased CPP was carried out using three-chambered CPP Med Associates boxes and software. The two end chambers have distinct visual (gray *vs* striped walls) and tactile (small grid *vs* large grid flooring) cues to allow differentiation. On the pre-test day, animals were allowed to freely explore all three chambers for 20 min. Groups were then attributed and pairing sides were adjusted to balance out any pre-existing chamber bias. Next, drug-context learning was achieved by pairing an injection of saline with one chamber in the morning, and a second injection of cocaine or morphine with the other chamber in the afternoon for two consecutive days. CPP testing was carried out on the fourth day when each animal was again allowed to explore all chambers freely. Place preference score was taken as the difference in time spent on the cocaine-paired side time *vs* on the saline-paired side.

### Self-administration and behavioral economics testing

Rats were anesthetized with ketamine (100 mg/kg) and xylazine (10 mg/kg) and implanted with chronic indwelling jugular catheters as described previously ^44^. Animals were singly housed, and all sessions took place during the active/dark cycle (12:00–15:00). After a 4-day recovery period, animals underwent training for self-administration where they were given access to a cocaine-paired lever on a fixed ratio one (FR1) schedule using a cocaine dose of 0.8 mg/kg/infusion delivered over 5 s. After each infusion, the lever was retracted and a stimulus light was illuminated for a 20 s timeout period. Responding on a second inactive lever was recorded but resulted in no programmed consequence. Acquisition was defined as the first session in which an animal allocated > 70% of their responses to the active lever, and when a stable pattern of inter-infusion intervals was present. After acquisition, rats then went through a within-session threshold procedure, described in detail below, before being split into two unbiased groups (Extended Data Fig. S5) for HSV surgery. After surgery and one recovery day, rats were subjected to 2 FR1 consumption sessions and 2 threshold sessions. The threshold procedure has been used by our lab ^44^ and others ^45, 46, 48–50^ to determine differences in consummatory and appetitive responding. Briefly, a descending series of 11-unit doses of cocaine (421, 237, 133, 75, 41, 24, 13, 7.5, 4.1, 2.4, and 1.3 µg/infusion) are available for 10 min upon an FR1 schedule with no timeouts, and levers remain extended. Consequently, the “price” of cocaine – the number of responses required to obtain 1 mg – increases over time. Both the number of lever presses and the total cocaine intake are measured for each price bin and plotted as a function of dose/price. The resulting demand curves can be used for mathematical curve fitting using the following equation: log_10_(*Q*) = log_10_(*Q*_0_) + *k* × (*e*^−α×*Q*_0_×*C*^ − 1) where *Q* is consumption and *C* is the varying cost of the reinforcer ^46, 48^. Here, curve fitting was achieved using custom-made R code utilizing the *stats::nls* function, and goodness of fit was assessed by computing a pseudo-R^2^ coefficient (the square of the correlation between observed and predicted log_10_(*Q*) values) and the Akaike Information Criterion (AIC) for each individual fit. The *k* value was determined by fitting all individual curves with 0.1 increments of *k* values between 1 and 5, and the *k* value that maximized the overall pseudo-R^2^ and minimized the overall AIC was selected and was here equal to 2.7.

The parameters Q_0_, Q_max_, P_max_, O_max_ and α were then calculated for each animal. Q_0_ is a measure of the preferred level of cocaine consumption of the animals and was measured as the amount of consumption at a theoretical minimally constraining price. P_max_ is the inflection point of the demand curve and was calculated as the point for which the first derivative slope of the demand function is equal to −1 in the log-log space, and corresponds to the price at which the animal no longer emits enough responses to maintain intake and consumption decreases. Q_max_ is the consumption at P_max_, and O_max_ the behavioral output (active lever presses) at P_max_. The measure of elasticity α, also termed the essential value, is a measure of how sensitive the demand is to price. With high elasticity, responding will drop off quickly as price increases, whereas with low elasticity, animals will be motivated to continue consuming drug, regardless of cost. Curve fitting and analysis code is available upon request.

### Statistics

No statistical power estimation analyses were used to predetermine sample sizes, which instead were chosen to match previous publications ^11, 31, 37, 44^. All statistics were performed in R v 4.0.2. Detailed statistics, including the exact functions and arguments used, are provided for each figure panel in Extended Data Table S6. In summary, pairwise comparisons were performed with Welch’s *t*-tests (*stats::t.test* function), enrichment tests using Fisher’s LSD (*stats::fisher.test* function), correlations using Pearson’s *r* (*stats::cor.test* function) and more complex multifactorial designs were analyzed using linear models computed with the *stats::lm* function for fixed effects-only models or *lmerTest::lmer* function for mixed effects models. Random effects (repeated measures and/or nested observations) were modeled as random intercept factors. Subsequent analysis of variance was performed using type III sums of squares with Kenward-Roger’s approximation of degrees of freedom. Of note, biological sex was always included as a fixed effect factor according to best practice guidelines in considering sex as a biological variable ^61^. Pooled data are represented in figures, but individual data points are color-coded by sex. *Post hoc* testing (on pooled data when relevant) was performed using the *emmeans* package and significance was adjusted using Sidak’s correction, except for RNAseq data where Benjamini-Hochberg correction was used as part of the *DESeq2* pipeline. Bar and line graphs represent mean ± sem. Correlation graphs represent regression line with its 95% confidence interval. Significance was set at *p* < 0.05.

## DATA AVAILABILITY

All RNAseq data reported in this study are deposited publicly in the Gene Expression Omnibus under accession code GSE198527. Unmodified Western blots scans are provided as Source Data. Other supporting raw data are available from the corresponding author upon request.

## CODE AVAILABILITY

Scripts and code utilized in this study, including for statistical analysis, are available from the corresponding author upon request.

## ETHICS DECLARATION

The authors declare no competing financial interests.

## ACKNOWLEDGEMENTS

This work was supported by funding from the Boehringer Ingelheim Fonds (PhD Fellowship to A.G.) and the National Institute of Health (P01DA047233 to E.J.N.). The authors would like to thank Katherine Beach, Catherine McManus, Kyra Schmidt and Ezekiell Mouzon for transgenics breeding and genotyping, Dr. Logan Brown and Dr. Boris Kantor from the Duke University Viral Vector Core for cloning and packaging AAV vectors, Dr. Guillermo Villegas and Dr. Edgardo Aritzia from the Dean’s Flow Cytometry CoRE at the Icahn School of Medicine at Mount Sinai for assistance in nuclei sorting, and Dr. Mark Baxter for statistical advice.

---

**Fig. S1.**
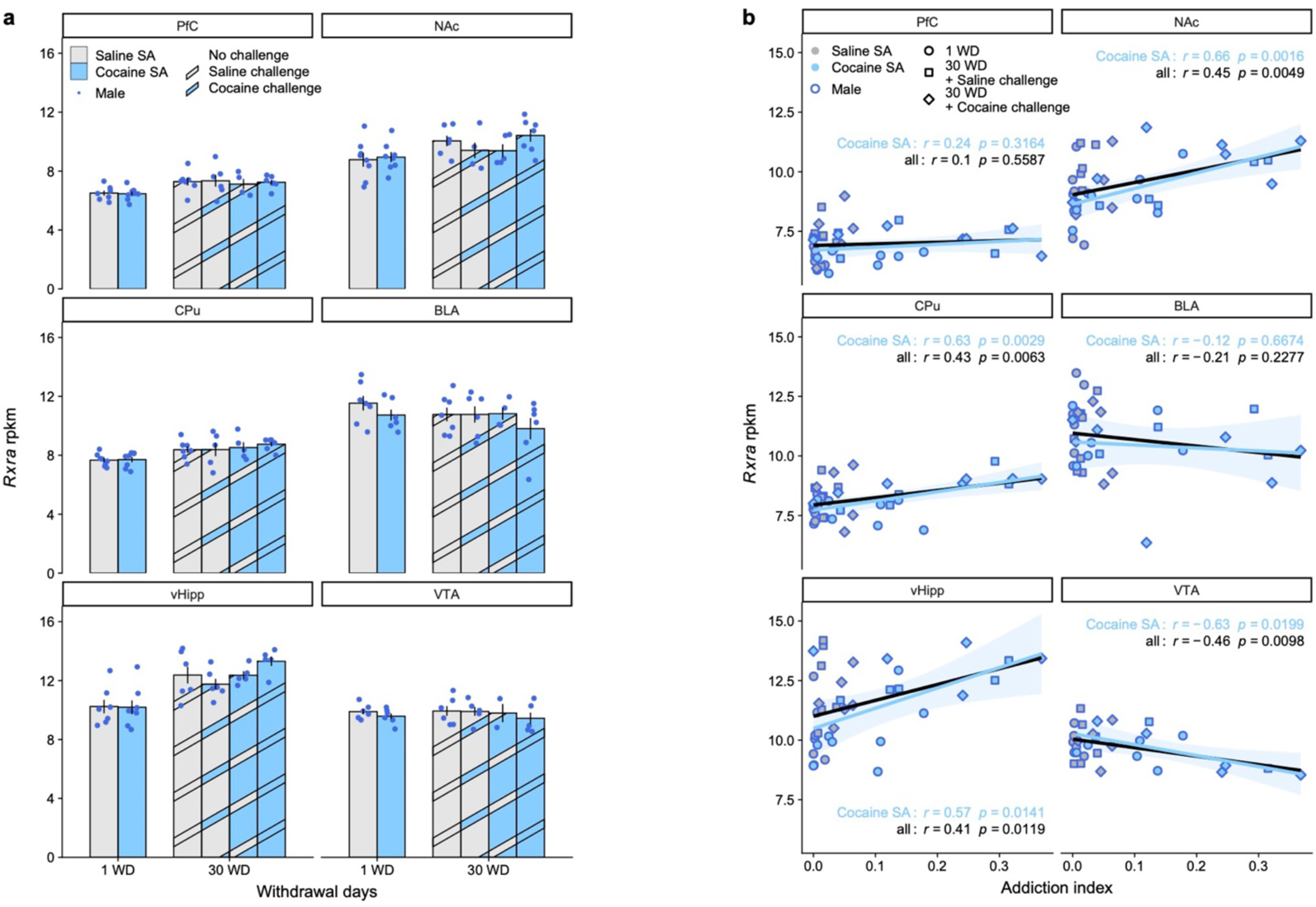
Brain-wide RXRα expression after cocaine self-administration. **a**, RNAseq of bulk NAc tissue showed no regulation of *Rxra* transcripts across experimental groups in any of the six brain regions analyzed (data from original study ^11^). **b**, Correlation between *Rxra* levels and the Addiction Index, a composite metric representative of addiction-like behavioral domains computed using exploratory factor analysis ^11^ was positive in NAc (across all samples: Pearson’s *r* = 0.45, *p* = 0.0049; across cocaine SA samples only: Pearson’s *r* = 0.66, *p* = 0.0016), CPu (across all samples: Pearson’s *r* = 0.43, *p* = 0.0063; across cocaine SA samples only: Pearson’s *r* = 0.63, *p* = 0.0029) and vHipp (across all samples: Pearson’s *r* = 0.41, *p* = 0.0119; across cocaine SA samples only: Pearson’s *r* = 0.57, *p* = 0.0141) and negative in VTA (across all samples: Pearson’s *r* = −0.46, *p* = 0.0098; across cocaine SA samples only: Pearson’s *r* = −0.63, *p* = 0.0199). Bar graphs represent mean ± sem. Correlation graphs represent regression line with its 95% confidence interval.

**Fig. S2.**
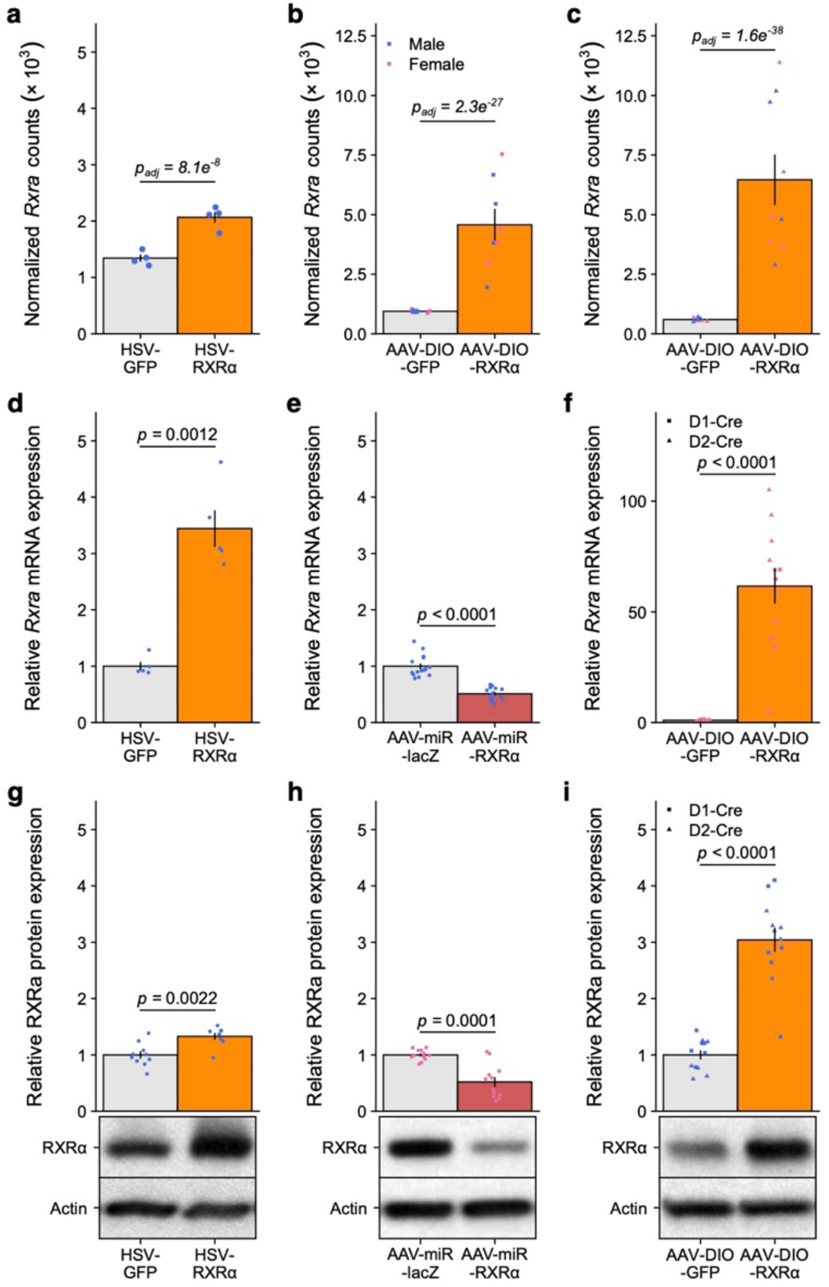
Validation of viral constructs. **a**, NAc *Rxra* transcripts count was significantly increased in male wild type mice injected with HSV-RXRα via RNAseq (Fig. 1 dataset; Wald’s test, *p_adj_* = 8.1e^−8^). **b**, *Rxra* transcripts count in D1-MSNs was significantly increased in male and female D1-Cre x ^fl/fl^eGFP::L10a mice injected with AAV-DIO-RXRα via RNAseq after FAN-sorting (Fig. 3 dataset; Wald’s test, *p_adj_* = 2.3e^−27^). **c**, *Rxra* transcripts count in D2-MSNs was significantly increased in male and female D2-Cre x ^fl/fl^eGFP::L10a mice injected with AAV-DIO-RXRα via RNAseq after FAN-sorting (Fig. 4 dataset; Wald’s test, *p_adj_* = 1.6e^−38^). **d**, NAc *Rxra* transcripts levels were significantly increased in male wild type mice injected with HSV-RXRα via RT-qPCR (unpaired Welch’s *t*-test: *t*_4.4178_ = −7.3332, *p* = 0.00124). **e**, NAc *Rxra* transcripts levels were significantly decreased in male wild type mice injected with AAV-miR-RXRα via RT-qPCR (unpaired Welch’s *t*-test: *t*_22.58_ = 8.4543, *p* = 1.899e^−8^). **f**, NAc *Rxra* transcripts levels were significantly increased in female D1-Cre and D2-Cre mice injected with AAV-DIO-RXRα via RT-qPCR (LMM-ANOVA: main effect of Virus: F_1,20_ = 77.2889, *p* = 1.559e^−8^, followed by Sidak’s *post hoc* tests). **g**, NAc RXRα protein levels were significantly increased in male wild type mice injected with HSV-RXRα via Western Blot (unpaired Welch’s *t*-test: *t*_15.847_ = 3.6574, *p* = 0.002155). **h**, NAc RXRα protein levels were significantly decreased in female wild type mice injected with AAV-miR-RXRα via Western Blot (unpaired Welch’s *t*-test: *t*_13.186_ = 5.2867, *p* = 0.0001404). **f**, NAc RXRα protein levels were significantly decreased in male D1-Cre and D2-Cre mice injected with AAV-DIO-RXRα via Western Blot (LMM-ANOVA: main effect of Virus: F_1,20_ = 74.8156, *p* = 3.418e^−8^, followed by Sidak’s *post hoc* tests). Representative Western Blot pictures for RXRα and control actin bands are attached to the corresponding quantification. Bar graphs represent mean ± sem.

**Fig. S3.**
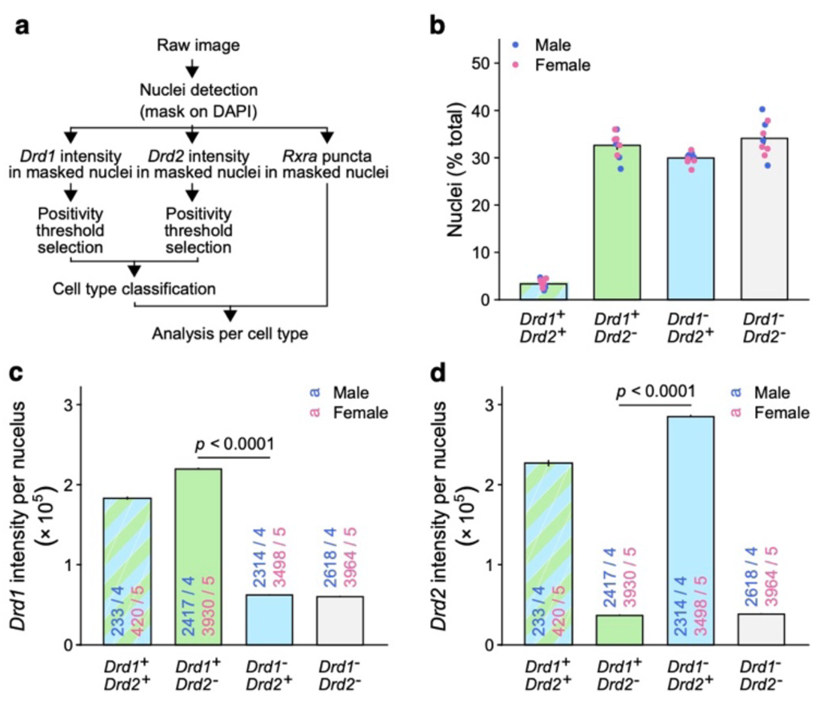
RNA FISH-based classification of NAc cell types. **a**, Schematic pipeline for RNA FISH image analysis, cell type classification and signal quantification. **b**, Detected and classified cell types recapitulated expected relative abundance in mouse NAc (LMM-ANOVA: main effect of CellType: F_3,28_ = 291.6805, *p* < 2e^− 16^, followed by Sidak’s *post hoc* tests). **c**, *Drd1* expression across NAc cell types (LMM-ANOVA: main effect of CellType: F_3,19381.2_ = 16327.88, *p* < 2e^−16^, followed by Sidak’s *post hoc* tests). **d**, *Drd2* expression across NAc cell types (LMM-ANOVA: main effect of CellType: F_3,19380.4_ = 21539.97, *p* < 2e^−16^, followed by Sidak’s *post hoc* tests). Bar graphs represent mean ± sem after combining male and female data.

**Fig. S4.**
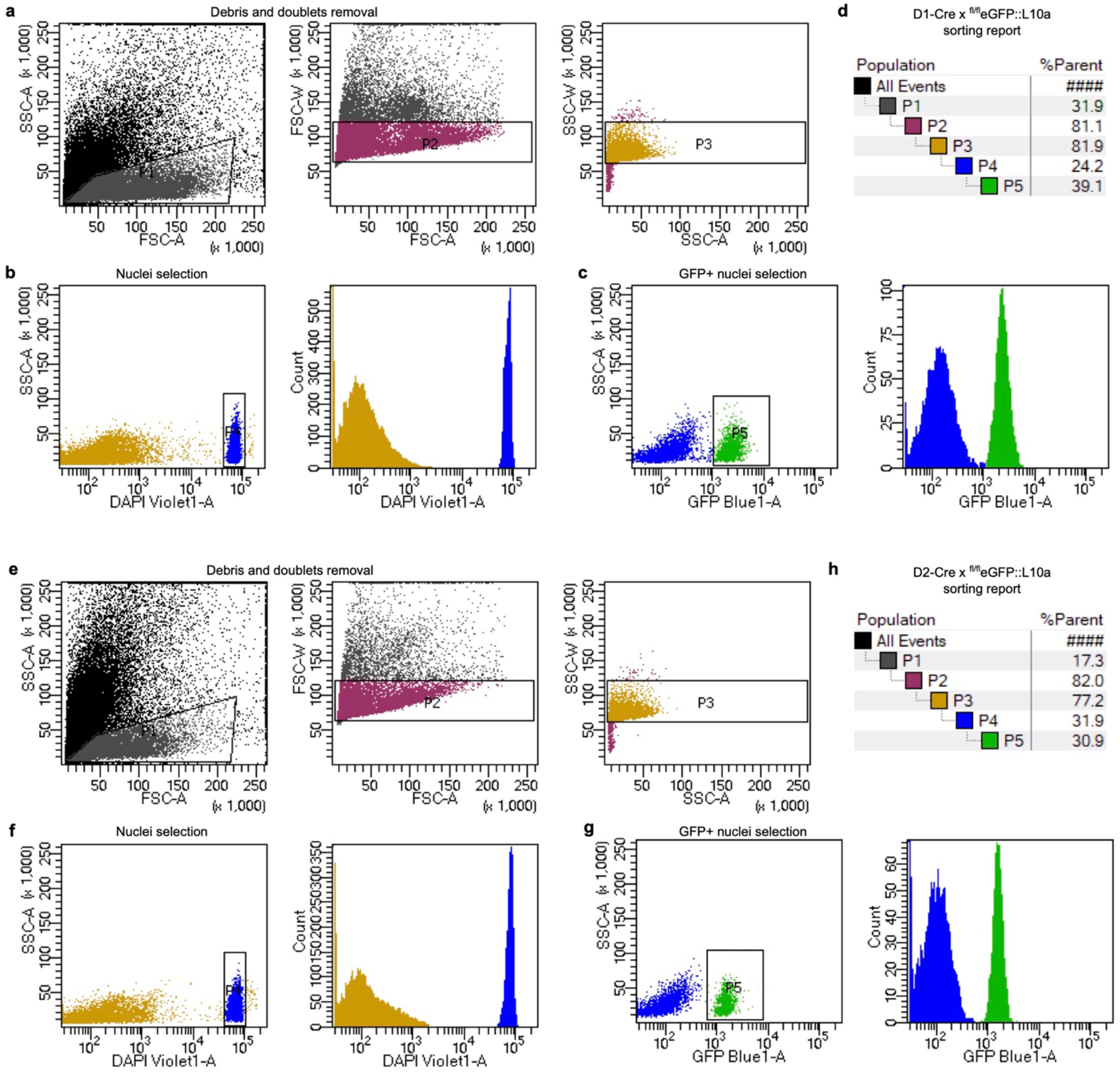
FAN-sorting of nuclei from D1- and D2-MSNs. **a**, Representative FANS gating strategy to exclude debris and doublets from a D1-Cre x ^fl/fl^eGFP::L10a sample. **b**, Representative FANS gating strategy to select nuclei (*ie* DAPI+ events) from a D1-Cre x ^fl/fl^eGFP::L10a sample. **c**, Representative FANS gating strategy to select D1-MSNs nuclei (*ie* GFP+ events) from a D1-Cre x ^fl/fl^eGFP::L10a sample. **c**, Summary of FAN-sorting hierarchical gating strategy for a D1-Cre x ^fl/fl^eGFP::L10a sample. **e**, Representative FANS gating strategy to exclude debris and doublets from a D2-Cre x ^fl/fl^eGFP::L10a sample. **f**, Representative FANS gating strategy to select nuclei (*ie* DAPI+ events) from a D2-Cre x ^fl/fl^eGFP::L10a sample. **g**, Representative FANS gating strategy to select D2-MSNs nuclei (*ie* GFP+ events) from a D2-Cre x ^fl/fl^eGFP::L10a sample. **h** Summary of FAN-sorting hierarchical gating strategy for a D2-Cre x ^fl/fl^eGFP::L10a sample. FSC = Forward Scatter, SSC = Side Scatter.

**Fig. S5.**
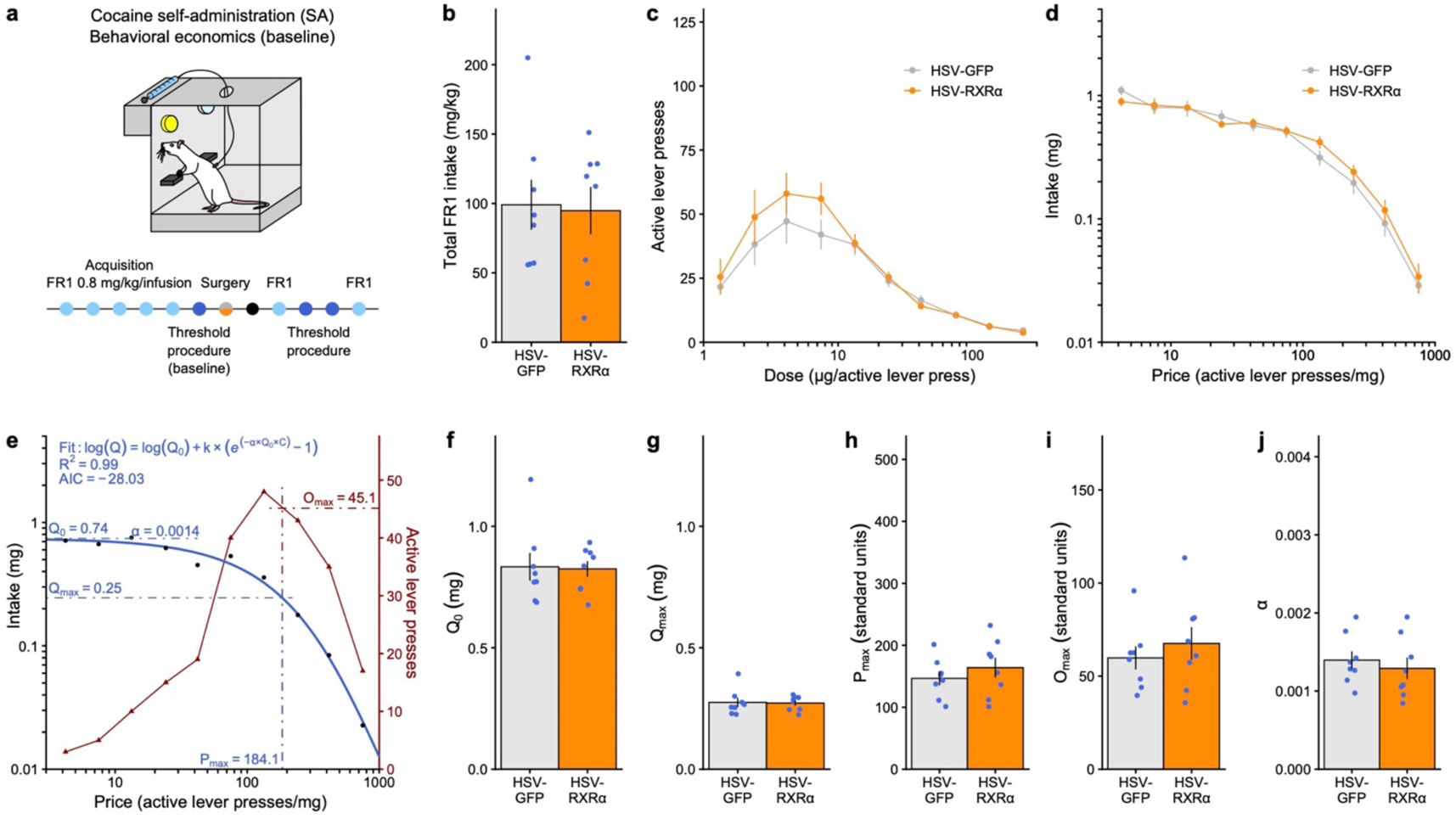
Behavioral economics metrics before RXRα overexpression. **a**, Experimental design for behavioral economics testing. Male rats (n = 8/group) were trained to self-administer cocaine (0.8 mg/kg/infusion) on a FR1 schedule of reinforcement before being injected in NAc with an HSV overexpressing RXRα or a control GFP sequence. Before surgery, animals went through the threshold procedure to assess motivation at baseline. **b**, Total cocaine intake in acquisition FR1 sessions was similar before group assignment (unpaired Welch’s *t*-test: *t*_13.961_ = 0.16909, *p* = 0.8682). **c**, Dose response curves in the threshold procedure showed similar responding before group assignment (LMM-ANOVA: interaction Dose:Virus: F_9,126_ = 0.7509, *p* = 0.6617, followed by Sidak’s *post hoc* tests). **d**, Averaged demand curves were also similar before group assignment (LMM-ANOVA: interaction Price:Virus: F_9,126_ = 0.9386, *p* = 0.4943, followed by Sidak’s *post hoc* tests). **e**, Representative example of task performance in the threshold procedure at baseline (from a rat later injected with HSV-GFP control), highlighting mathematical demand curve fitting and extraction of behavioral economics parameters. Goodness of fit was assessed by computing a pseudo-R^2^ coefficient and the Akaike Information Criterion (AIC) for each individual fit. **f**, Consumption at low effort Q_0_ (unpaired Welch’s *t*-test: *t*_11.097_ = 0.12951, *p* = 0.8993), **g**, consumption at maximum effort Q_max_ (unpaired Welch’s *t*-test: *t*_11.046_ = 0.12078, *p* = 0.906), motivation metrics **h**, P_max_ (unpaired Welch’s *t*-test: *t*_12.516_ = −0.8658, *p* = 0.4029) and **i**, O_max_ (unpaired Welch’s *t*-test: *t*_12.594_ = −0.7262, *p* = 0.481), and **j**, demand elasticity α (unpaired Welch’s *t*-test: *t*_13.428_ = 0.59101, *p* = 0.5643), were similar before group assignment. Bar graphs and line graphs represent mean ± sem.

