## Supplemental Table S6 for "Transcriptional control of nucleus accumbens neuronal excitability by Retinoid X Receptor Alpha tunes sensitivity to drug rewards"

| Figure | Statistics input and output |
| --- | --- |
| Fig. 1b | Call: voomlimma pairwise comparisons ^13^ Region \| Pairwise \| logFC \| AveExpr \| t \| P.Value \| adj.P.Val \| B; NAc \| CNvSN \| 0.03696451 \| 5.69822834 \| 0.41106822 \| 0.68332724 \| 0.99988234 \| -5.0343219; NAc \| CSvSS \| -0.0981674 \| 5.69822834 \| -0.9116555 \| 0.36768669 \| 0.98608515 \| -4.7566445; NAc \| SCvSS \| -0.0291495 \| 5.69822834 \| -0.2720538 \| 0.78705062 \| 0.99999207 \| -4.9654173; NAc \| CCvSS \| 0.04840867 \| 5.69822834 \| 0.49075105 \| 0.62641938 \| 0.99967096 \| -4.960794 |
| Fig. 1c | Call: stats::cor.test(Rxra.rpkm, Addiction.Index, method = “pearson”)  *All samples*  Pearson's product-moment correlation  data: Rxra.rpkm and Addiction.Index  t = 3.0017, df = 36, p-value = 0.004855  alternative hypothesis: true correlation is not equal to 0  95 percent confidence interval:  0.1490520 0.6711097  sample estimates:  cor  0.4474162  *Cocaine samples only*  Pearson's product-moment correlation  data: Rxra.rpkm and Addiction.Index  t = 3.7087, df = 18, p-value = 0.001607  alternative hypothesis: true correlation is not equal to 0  95 percent confidence interval:  0.3042280 0.8524079  sample estimates:  cor  0.6581446 |
| Fig. 1e | Call: stats::fisher.test(contmat, alternative = “greater”)  Fisher's Exact Test for Count Data  data: contmat  p-value < 2.2e-16  alternative hypothesis: true odds ratio is greater than 1  95 percent confidence interval:  3.009465 Inf  sample estimates:  odds ratio  3.8235 |
| Fig. 2b | Call: lm(norm.expr ~ Drug*Time*Sex) %>% car::Anova(type = “III”, test.statistic = “F)  Anova Table (Type III tests)  Response: norm.expr  Sum Sq Df F value Pr(>F)  (Intercept) 42.939 1 1176.8535 <0.0000000000000002 ***  Drug 0.030 1 0.8185 0.3726  Time 0.012 1 0.3323 0.5685  Sex 0.000 1 0.0010 0.9745  Drug:Time 0.094 1 2.5762 0.1186  Drug:Sex 0.016 1 0.4472 0.5086  Time:Sex 0.057 1 1.5560 0.2216  Drug:Time:Sex 0.000 1 0.0056 0.9410  Residuals 1.131 31  ---  Signif. codes: 0 ‘***’ 0.001 ‘**’ 0.01 ‘*’ 0.05 ‘.’ 0.1 ‘ ’ 1  Posthoc: emmeans(model, c(“Drug”, “Time”)) %>% pairs(simple=”each”, adjust = “sidak”)  Time Drug contrast estimate SE df t.ratio p.value  30 . Saline - Cocaine -0.1540 0.0881 31 -1.748 0.3151  60 . Saline - Cocaine 0.0430 0.0854 31 0.503 0.9788  . Saline 30 - 60 -0.1338 0.0881 31 -1.520 0.4497  . Cocaine 30 - 60 0.0631 0.0854 31 0.739 0.9185  Results are averaged over some or all of the levels of: Sex  P value adjustment: sidak method for 4 tests |
| Fig. 2c | Call: lm(norm.expr ~ Drug*Time*Sex) %>% car::Anova(type = “III”, test.statistic = “F)  Anova Table (Type III tests)  Response: norm.expr  Sum Sq Df F value Pr(>F)  (Intercept) 32.971 1 852.9357 < 0.00000000000000022 ***  Drug 0.004 1 0.1004 0.7534207  Time 0.611 1 15.8177 0.0003734 ***  Sex 0.256 1 6.6347 0.0148239 *  Drug:Time 0.019 1 0.4864 0.4905546  Drug:Sex 0.011 1 0.2862 0.5963335  Time:Sex 0.001 1 0.0345 0.8537341  Drug:Time:Sex 0.192 1 4.9578 0.0331360 *  Residuals 1.237 32  ---  Signif. codes: 0 ‘***’ 0.001 ‘**’ 0.01 ‘*’ 0.05 ‘.’ 0.1 ‘ ’ 1  Posthoc: emmeans(model, c(“Drug”, “Time”)) %>% pairs(simple=”each”, adjust = “sidak”)  Time Drug contrast estimate SE df t.ratio p.value  30 . Saline - Cocaine -0.0631 0.0879 32 -0.717 0.9260  60 . Saline - Cocaine 0.0237 0.0879 32 0.269 0.9980  . Saline 30 - 60 0.2039 0.0879 32 2.319 0.1035  . Cocaine 30 - 60 0.2906 0.0879 32 3.305 0.0093  Results are averaged over some or all of the levels of: Sex  P value adjustment: sidak method for 4 tests |
| Fig. 2d | Call: lm(norm.expr ~ Drug*Time*Sex) %>% car::Anova(type = “III”, test.statistic = “F)  Anova Table (Type III tests)  Response: norm.expr  Sum Sq Df F value Pr(>F)  (Intercept) 43.657 1 899.7109 < 0.0000000000000002 ***  Drug 0.071 1 1.4631 0.23530  Time 0.000 1 0.0001 0.99260  Sex 0.004 1 0.0843 0.77348  Drug:Time 0.000 1 0.0070 0.93398  Drug:Sex 0.164 1 3.3832 0.07516 .  Time:Sex 0.000 1 0.0002 0.98944  Drug:Time:Sex 0.114 1 2.3585 0.13443  Residuals 1.553 32  ---  Signif. codes: 0 ‘***’ 0.001 ‘**’ 0.01 ‘*’ 0.05 ‘.’ 0.1 ‘ ’ 1  Posthoc: emmeans(model, c(“Drug”, “Time”)) %>% pairs(simple=”each”, adjust = “sidak”)  Time Drug contrast estimate SE df t.ratio p.value  30 . Saline - Cocaine -0.09007 0.0985 32 -0.914 0.8398  60 . Saline - Cocaine -0.07844 0.0985 32 -0.796 0.8957  . Saline 30 - 60 -0.00516 0.0985 32 -0.052 1.0000  . Cocaine 30 - 60 0.00647 0.0985 32 0.066 1.0000  Results are averaged over some or all of the levels of: Sex  P value adjustment: sidak method for 4 tests |
| Fig. 2e mRNA | Call: stats::cor.test(Target, Rxra, method = “pearson”) \| Saline \| cor; Arc \| t = 3.0322, df = 17, p-value = 0.007519 \| 0.5924597; cFos \| t = 2.0837, df = 17, p-value = 0.05259 \| 0.4510388; FosB \| t = 4.4898, df = 17, p-value = 0.0003227 \| 0.7365476; deltaFosB \| t = 3.517, df = 17, p-value = 0.002645 \| 0.6489703; Homer1a \| t = 1.8417, df = 17, p-value = 0.08303 \| 0.4078453; Npas4 \| t = 2.4958, df = 17, p-value = 0.02315 \| 0.5178395; Nr4a1 \| t = 2.9073, df = 17, p-value = 0.00981 \| 0.5762681; Zif268 \| t = 6.0513, df = 17, p-value = 0.00001296 \| 0.8264026; Drd1 \| t = 7.2938, df = 17, p-value = 0.000001255 \| 0.8705359; Drd2 \| t = 3.1824, df = 17, p-value = 0.00545 \| 0.6110075; Gria1 \| t = 7.6353, df = 17, p-value = 0.0000006849 \| 0.879902; Grin1 \| t = 8.0754, df = 17, p-value = 0.0000003211 \| 0.8906284; \| Cocaine \| cor; Arc \| t = 2.5278, df = 18, p-value = 0.02105 \| 0.5118425; cFos \| t = 1.5572, df = 18, p-value = 0.1368 \| 0.3445555; FosB \| t = 2.6276, df = 18, p-value = 0.01708 \| 0.5265328; deltaFosB \| t = 1.94, df = 18, p-value = 0.06822 \| 0.4158429; Homer1a \| t = 1.4347, df = 18, p-value = 0.1685 \| 0.3203495; Npas4 \| t = 0.88881, df = 18, p-value = 0.3858 \| 0.2050423; Nr4a1 \| t = 2.6142, df = 18, p-value = 0.01757 \| 0.5245843; Zif268 \| t = 4.5718, df = 18, p-value = 0.0002365 \| 0.7329993; Drd1 \| t = 5.4541, df = 18, p-value = 0.00003512 \| 0.7893137; Drd2 \| t = 3.5688, df = 18, p-value = 0.002194 \| 0.643723; Gria1 \| t = 5.344, df = 18, p-value = 0.00004437 \| 0.7831912; Grin1 \| t = 5.2626, df = 18, p-value = 0.00005278 \| 0.7785134; \| All samples \| cor; Arc \| t = 3.8494, df = 37, p-value = 0.0004531 \| 0.5347488; cFos \| t = 2.3018, df = 37, p-value = 0.02707 \| 0.3539252; FosB \| t = 4.4042, df = 37, p-value = 0.00008728 \| 0.586464; deltaFosB \| t = 3.1416, df = 37, p-value = 0.0033 \| 0.4588893; Homer1a \| t = 2.6151, df = 37, p-value = 0.01283 \| 0.3949677; Npas4 \| t = 2.7979, df = 37, p-value = 0.008118 \| 0.4178835; Nr4a1 \| t = 3.7911, df = 37, p-value = 0.0005366 \| 0.5289284; Zif268 \| t = 6.517, df = 37, p-value = 0.0000001258 \| 0.7310414; Drd1 \| t = 9.6225, df = 37, p-value = 0.00000000001296 \| 0.8452739; Drd2 \| t = 4.9505, df = 37, p-value = 0.00001644 \| 0.6312244; Gria1 \| t = 9.5636, df = 37, p-value = 0.00000000001526 \| 0.8437888; Grin1 \| t = 9.8511, df = 37, p-value = 0.000000000006895 \| 0.8508656 |
| Fig. 2e  Protein | Call: stats::cor.test(Target, Rxra, method = “pearson”) \| Saline \| cor; Arc \| t = 0.31064, df = 18, p-value = 0.7596 \| 0.07302229; pCREB \| t = 4.4916, df = 18, p-value = 0.0002822 \| 0.72697; pERK \| t = 1.5146, df = 18, p-value = 0.1472 \| 0.3362167; GluN2B \| t = 0.14213, df = 18, p-value = 0.8886 \| 0.03348223; PSD95 \| t = 1.7643, df = 18, p-value = 0.09464 \| 0.3839815; \| Cocaine \| cor; Arc \| t = 2.8544, df = 18, p-value = 0.01053 \| 0.5582053; pCREB \| t = 1.8265, df = 18, p-value = 0.0844 \| 0.3954273; pERK \| t = -0.56266, df = 18, p-value = 0.5806 \| -0.1314689; GluN2B \| t = 2.439, df = 18, p-value = 0.02531 \| 0.4983902; PSD95 \| t = -0.056814, df = 18, p-value = 0.9553 \| -0.01339; \| All samples \| cor; Arc \| t = 2.3238, df = 38, p-value = 0.02558 \| 0.3527355; pCREB \| t = 3.6952, df = 38, p-value = 0.0006896 \| 0.5141442; pERK \| t = 0.45839, df = 38, p-value = 0.6493 \| 0.07415578; GluN2B \| t = 1.5157, df = 38, p-value = 0.1379 \| 0.2387685; PSD95 \| t = 1.2074, df = 38, p-value = 0.2347 \| 0.1922109 |
| Fig. 2e  Nuclear  protein | Call: stats::cor.test(Target, Rxra, method = “pearson”) \| Saline \| cor; pCREB \| t = 1.2859, df = 18, p-value = 0.2148 \| 0.2900565; pERK \| t = 2.2833, df = 18, p-value = 0.03478 \| 0.4739039; p65/RelA \| t = 0.61405, df = 18, p-value = 0.5469 \| 0.1432396; \| Cocaine \| cor; pCREB \| t = 2.7574, df = 18, p-value = 0.01297 \| 0.5449413; pERK \| t = 3.1044, df = 18, p-value = 0.00612 \| 0.5905163; p65/RelA \| t = 2.7276, df = 18, p-value = 0.01382 \| 0.5407827; \| All samples \| cor; pCREB \| t = 2.9041, df = 38, p-value = 0.006106 \| 0.4261763; pERK \| t = 3.8227, df = 38, p-value = 0.0004761 \| 0.5270147; p65/RelA \| t = 2.3048, df = 38, p-value = 0.02673 \| 0.3502115 |
| Fig. 2f | Call: stats::cor.test(Zif268, Rxra, method = “pearson”)  *All samples*  Pearson's product-moment correlation  data: Zif268 and Rxra  t = 6.517, df = 37, p-value = 0.0000001258  alternative hypothesis: true correlation is not equal to 0  95 percent confidence interval:  0.5401021 0.8504069  sample estimates:  cor  0.7310414  *Cocaine samples only*  Pearson's product-moment correlation  data: Zif268 and Rxra  t = 4.5718, df = 18, p-value = 0.0002365  alternative hypothesis: true correlation is not equal to 0  95 percent confidence interval:  0.4299356 0.8876087  sample estimates:  cor  0.7329993  *Saline samples only*  Pearson's product-moment correlation  data: Zif268 and Rxra  t = 6.0513, df = 17, p-value = 0.00001296  alternative hypothesis: true correlation is not equal to 0  95 percent confidence interval:  0.5958522 0.9311105  sample estimates:  cor  0.8264026 |
| Fig. 2g | Call: stats::cor.test(pCREB, RXRa, method = “pearson”)  *All samples*  Pearson's product-moment correlation  data: pCREB and RXRa  t = 3.6952, df = 38, p-value = 0.0006896  alternative hypothesis: true correlation is not equal to 0  95 percent confidence interval:  0.2412782 0.7116717  sample estimates:  cor  0.5141442  *Cocaine samples only*  Pearson's product-moment correlation  data: pCREB and RXRa  t = 1.8265, df = 18, p-value = 0.0844  alternative hypothesis: true correlation is not equal to 0  95 percent confidence interval:  -0.05708198 0.71315652  sample estimates:  cor  0.3954273  *Saline samples only*  Pearson's product-moment correlation  data: pCREB and RXRa  t = 4.4916, df = 18, p-value = 0.0002822  alternative hypothesis: true correlation is not equal to 0  95 percent confidence interval:  0.4193557 0.8848388  sample estimates:  cor  0.72697 |
| Fig. 2h | Call: stats::cor.test(pCREB.P1, RXRa.P1, method = “pearson”)  *All samples*  Pearson's product-moment correlation  data: pCREB.P1 and RXRa.P1  t = 2.9041, df = 38, p-value = 0.006106  alternative hypothesis: true correlation is not equal to 0  95 percent confidence interval:  0.1322202 0.6512298  sample estimates:  cor  0.4261763  *Cocaine samples only*  Pearson's product-moment correlation  data: pCREB.P1 and RXRa.P1  t = 2.7574, df = 18, p-value = 0.01297  alternative hypothesis: true correlation is not equal to 0  95 percent confidence interval:  0.1349677 0.7956039  sample estimates:  cor  0.5449413  *Saline samples only*  Pearson's product-moment correlation  data: pCREB.P1 and RXRa.P1  t = 1.2859, df = 18, p-value = 0.2148  alternative hypothesis: true correlation is not equal to 0  95 percent confidence interval:  -0.1749158 0.6492430  sample estimates:  cor  0.2900565 |
| Fig. 3b | Call: lmerTest::lmer(Rxra.puncta ~Cell.type*Sex + (1\|mouseid) %>% anova(")  Type III Analysis of Variance Table with Satterthwaite's method  Sum Sq Mean Sq NumDF DenDF F value Pr(>F)  Sex 1.67 1.672 1 10 0.7011 0.4220120  Cell.type 323.04 107.680 3 19381 45.1531 < 2.2e-16 ***  Sex:Cell.type 41.87 13.958 3 19381 5.8530 0.0005442 ***  ---  Signif. codes: 0 ‘***’ 0.001 ‘**’ 0.01 ‘*’ 0.05 ‘.’ 0.1 ‘ ’ 1  Posthoc: emmeans(model, c(“Cell.type”)) %>% pairs(simple=”each”, adjust = “sidak”)  contrast estimate SE df z.ratio p.value  (Drd2-pos Drd1-pos) - (Drd2-neg Drd1-pos) 0.0531 0.0662 Inf 0.802 0.9630  (Drd2-pos Drd1-pos) - (Drd2-pos Drd1-neg) 0.3468 0.0665 Inf 5.219 <.0001  (Drd2-pos Drd1-pos) - (Drd2-neg Drd1-neg) 0.2990 0.0661 Inf 4.520 <.0001  (Drd2-neg Drd1-pos) - (Drd2-pos Drd1-neg) 0.2938 0.0288 Inf 10.214 <.0001  (Drd2-neg Drd1-pos) - (Drd2-neg Drd1-neg) 0.2459 0.0279 Inf 8.799 <.0001  (Drd2-pos Drd1-neg) - (Drd2-neg Drd1-neg) -0.0479 0.0284 Inf -1.684 0.4403  Results are averaged over some or all of the levels of: Sex  Degrees-of-freedom method: asymptotic  P value adjustment: sidak method for 6 tests |
| Fig. 3d | Call: DESeq2::DESeqDataSetFromMatrix(countData = GFP, colData = meta, design = ~ Sex + CellType) %>% DESeq2::DESeq() %>% DESeq2::results(contrast = c("CellType","D1","D2"), independentFiltering = T) mgi_symbol \| baseMean \| log2FoldChange \| lfcSE \| stat \| pvalue \| padj; Drd1 \| 626.790779 \| 2.02036602 \| 0.22696734 \| 8.90157141 \| 5.51E-19 \| 8.69E-16; Pdyn \| 452.384594 \| 1.76300937 \| 0.13033078 \| 13.5271908 \| 1.08E-41 \| 2.73E-38; Slc35d3 \| 94.721451 \| 1.52490011 \| 0.21456017 \| 7.10709795 \| 1.19E-12 \| 1.25E-09; Drd2 \| 6938.00707 \| -3.4760607 \| 0.08639985 \| -40.232254 \| 0 \| 0; Penk \| 4116.83416 \| -2.7136245 \| 0.30935207 \| -8.7719616 \| 1.76E-18 \| 2.61E-15; Adora2a \| 718.390073 \| -1.8159854 \| 0.16878725 \| -10.75902 \| 5.37E-27 \| 1.04E-23 |
| Fig. 3e | Call: DESeq2::DESeqDataSetFromMatrix(countData = GFP, colData = meta, design = ~ Sex + CellType) %>% DESeq2::DESeq() %>% DESeq2::results(contrast = c("CellType","D1","D2"), independentFiltering = T) mgi_symbol \| baseMean \| log2FoldChange \| lfcSE \| stat \| pvalue \| padj; Rxra \| 762.002154 \| 0.19339545 \| 0.06212004 \| 3.11325377 \| 0.00185037 \| 0.04999662 |
| Fig. 3f | Call: stats::fisher.test(contmat, alternative = “greater”)  Fisher's Exact Test for Count Data  data: contmat  p-value < 2.2e-16  alternative hypothesis: true odds ratio is greater than 1  95 percent confidence interval:  2.099507 Inf  sample estimates:  odds ratio  2.252595 |
| Fig. 3i | All statistics available in Table S2 and Table S3 |
| Fig. 4c | Call : lmerTest ::lmer(RMP ~ Virus*Sex*Cell.type + (1\|Mouse), control=lmerControl(check.nobs.vs.nRE="ignore", calc.derivs = F)) %>% anova(ddf = "Kenward-Roger")  Type III Analysis of Variance Table with Kenward-Roger's method  Sum Sq Mean Sq NumDF DenDF F value Pr(>F)  Virus 27.157 27.157 1 4.978 0.8204 0.4068  Sex 18.226 18.226 1 5.258 0.5506 0.4899  Cell.type 46.198 46.198 1 57.479 1.3955 0.2423  Virus:Sex 120.440 120.440 1 5.258 3.6383 0.1119  Virus:Cell.type 51.418 51.418 1 57.479 1.5532 0.2177  Sex:Cell.type 13.526 13.526 1 57.479 0.4086 0.5252  Virus:Sex:Cell.type 10.936 10.936 1 57.479 0.3304 0.5677  Posthoc: emmeans(model, c(“Virus”, “Cell.type”)) %>% pairs(simple=”each”, adjust = “sidak”)  Cell.type Virus contrast estimate SE df t.ratio p.value  D1-MSNs . (AAV-miR\n-lacZ) - (AAV-miR\n-RXRa) -3.0803 1.96 13.8 -1.570 0.4504  D2-MSNs . (AAV-miR\n-lacZ) - (AAV-miR\n-RXRa) 0.4784 2.09 21.8 0.229 0.9990  . AAV-miR\n-lacZ (D1-MSNs) - (D2-MSNs) -3.4659 2.00 55.0 -1.737 0.3083  . AAV-miR\n-RXRa (D1-MSNs) - (D2-MSNs) 0.0927 2.04 59.1 0.045 1.0000  Results are averaged over some or all of the levels of: Sex  Degrees-of-freedom method: kenward-roger  P value adjustment: sidak method for 4 tests |
| Fig. 4d | Call : lmerTest ::lmer(rheo ~ Virus*Sex*Cell.type + (1\|Mouse), control=lmerControl(check.nobs.vs.nRE="ignore", calc.derivs = F)) %>% anova(ddf = "Kenward-Roger")  Type III Analysis of Variance Table with Kenward-Roger's method  Sum Sq Mean Sq NumDF DenDF F value Pr(>F)  Virus 15.76 15.76 1 5.790 0.0207 0.8904  Sex 471.87 471.87 1 7.330 0.6211 0.4554  Cell.type 1696.02 1696.02 1 55.600 2.2325 0.1408  Virus:Sex 5.53 5.53 1 7.330 0.0073 0.9343  Virus:Cell.type 1564.60 1564.60 1 55.600 2.0595 0.1569  Sex:Cell.type 715.85 715.85 1 55.456 0.9423 0.3359  Virus:Sex:Cell.type 196.46 196.46 1 55.456 0.2586 0.6131  Posthoc: emmeans(model, c(“Virus”, “Cell.type”)) %>% pairs(simple=”each”, adjust = “sidak”)  Cell.type Virus contrast estimate SE df t.ratio p.value  D1-MSNs . (AAV-miR\n-lacZ) - (AAV-miR\n-RXRa) 12.495 18.32 7.69 0.682 0.9447  D2-MSNs . (AAV-miR\n-lacZ) - (AAV-miR\n-RXRa) -7.623 18.30 8.06 -0.417 0.9905  . AAV-miR\n-lacZ (D1-MSNs) - (D2-MSNs) -0.414 9.56 53.56 -0.043 1.0000  . AAV-miR\n-RXRa (D1-MSNs) - (D2-MSNs) -20.532 10.26 57.15 -2.002 0.1856  Results are averaged over some or all of the levels of: Sex  Degrees-of-freedom method: kenward-roger  P value adjustment: sidak method for 4 tests |
| Fig. 4f | Call: lmerTest::lmer(AP.number ~ Virus*Sex*Cell.type*Current + (1\|Mouse) + (1\|Neuron), control=lmerControl(check.nobs.vs.nRE="ignore", calc.derivs = F)) %>% anova(ddf = "Kenward-Roger")  Type III Analysis of Variance Table with Kenward-Roger's method  Sum Sq Mean Sq NumDF DenDF F value Pr(>F)  Virus 19.7 19.7 1 6.75 3.5393 0.103534  Sex 0.9 0.9 1 7.35 0.1591 0.701304  Cell.type 2.3 2.3 1 83.47 0.4118 0.522838  Current 17827.8 17827.8 1 1012.00 3205.6799 < 2.2e-16 ***  Virus:Sex 1.6 1.6 1 7.35 0.2915 0.605194  Virus:Cell.type 12.9 12.9 1 83.47 2.3174 0.131713  Sex:Cell.type 1.8 1.8 1 83.24 0.3306 0.566853  Virus:Current 1632.8 1632.8 1 1012.00 293.5920 < 2.2e-16 ***  Sex:Current 0.1 0.1 1 1012.00 0.0138 0.906543  Cell.type:Current 9.8 9.8 1 1012.00 1.7697 0.183719  Virus:Sex:Cell.type 2.8 2.8 1 83.24 0.5091 0.477514  Virus:Sex:Current 47.8 47.8 1 1012.00 8.5910 0.003454 **  Virus:Cell.type:Current 341.7 341.7 1 1012.00 61.4425 1.150e-14 ***  Sex:Cell.type:Current 14.9 14.9 1 1012.00 2.6720 0.102438  Virus:Sex:Cell.type:Current 91.5 91.5 1 1012.00 16.4517 5.374e-05 ***  ---  Signif. codes: 0 ‘***’ 0.001 ‘**’ 0.01 ‘*’ 0.05 ‘.’ 0.1 ‘ ’ 1  Posthoc: emmeans(model, c(“Current”, “Virus”, “Cell.type”)) %>% pairs(simple=”each”, adjust = “sidak”)  Cell.type Current contrast estimate SE df t.ratio p.value  D1-MSNs 0 (AAV-miR\n-lacZ) - (AAV-miR\n-RXRa) -0.0444 1.04 35.4 -0.043 1.0000  D2-MSNs 0 (AAV-miR\n-lacZ) - (AAV-miR\n-RXRa) 0.0290 1.10 52.4 0.026 1.0000  D1-MSNs 20 (AAV-miR\n-lacZ) - (AAV-miR\n-RXRa) -0.0444 1.04 35.4 -0.043 1.0000  D2-MSNs 20 (AAV-miR\n-lacZ) - (AAV-miR\n-RXRa) 0.0290 1.10 52.4 0.026 1.0000  D1-MSNs 40 (AAV-miR\n-lacZ) - (AAV-miR\n-RXRa) -0.0444 1.04 35.4 -0.043 1.0000  D2-MSNs 40 (AAV-miR\n-lacZ) - (AAV-miR\n-RXRa) 0.0290 1.10 52.4 0.026 1.0000  D1-MSNs 60 (AAV-miR\n-lacZ) - (AAV-miR\n-RXRa) -0.1278 1.04 35.4 -0.122 1.0000  D2-MSNs 60 (AAV-miR\n-lacZ) - (AAV-miR\n-RXRa) -0.0424 1.10 52.4 -0.039 1.0000  D1-MSNs 80 (AAV-miR\n-lacZ) - (AAV-miR\n-RXRa) 0.0121 1.04 35.4 0.012 1.0000  D2-MSNs 80 (AAV-miR\n-lacZ) - (AAV-miR\n-RXRa) 0.0220 1.10 52.4 0.020 1.0000  D1-MSNs 100 (AAV-miR\n-lacZ) - (AAV-miR\n-RXRa) 0.3008 1.04 35.4 0.288 1.0000  D2-MSNs 100 (AAV-miR\n-lacZ) - (AAV-miR\n-RXRa) 0.3425 1.10 52.4 0.311 1.0000  D1-MSNs 120 (AAV-miR\n-lacZ) - (AAV-miR\n-RXRa) 1.0079 1.04 35.4 0.966 1.0000  D2-MSNs 120 (AAV-miR\n-lacZ) - (AAV-miR\n-RXRa) 0.6193 1.10 52.4 0.562 1.0000  D1-MSNs 140 (AAV-miR\n-lacZ) - (AAV-miR\n-RXRa) 1.6044 1.04 35.4 1.537 0.9897  D2-MSNs 140 (AAV-miR\n-lacZ) - (AAV-miR\n-RXRa) 1.7830 1.10 52.4 1.619 0.9772  D1-MSNs 160 (AAV-miR\n-lacZ) - (AAV-miR\n-RXRa) 3.1353 1.04 35.4 3.004 0.1447  D2-MSNs 160 (AAV-miR\n-lacZ) - (AAV-miR\n-RXRa) 2.9883 1.10 52.4 2.714 0.2509  D1-MSNs 180 (AAV-miR\n-lacZ) - (AAV-miR\n-RXRa) 3.9204 1.04 35.4 3.756 0.0197  D2-MSNs 180 (AAV-miR\n-lacZ) - (AAV-miR\n-RXRa) 4.4278 1.10 52.4 4.021 0.0060  D1-MSNs 200 (AAV-miR\n-lacZ) - (AAV-miR\n-RXRa) 3.8484 1.04 35.4 3.687 0.0239  D2-MSNs 200 (AAV-miR\n-lacZ) - (AAV-miR\n-RXRa) 6.2334 1.10 52.4 5.661 <.0001  D1-MSNs 220 (AAV-miR\n-lacZ) - (AAV-miR\n-RXRa) 3.6139 1.04 35.4 3.462 0.0444  D2-MSNs 220 (AAV-miR\n-lacZ) - (AAV-miR\n-RXRa) 7.6609 1.10 52.4 6.957 <.0001  D1-MSNs 240 (AAV-miR\n-lacZ) - (AAV-miR\n-RXRa) 3.3216 1.04 35.4 3.182 0.0929  D2-MSNs 240 (AAV-miR\n-lacZ) - (AAV-miR\n-RXRa) 8.6580 1.10 52.4 7.862 <.0001  D1-MSNs 260 (AAV-miR\n-lacZ) - (AAV-miR\n-RXRa) 2.8252 1.04 35.4 2.707 0.2842  D2-MSNs 260 (AAV-miR\n-lacZ) - (AAV-miR\n-RXRa) 9.0419 1.10 52.4 8.211 <.0001  D1-MSNs 280 (AAV-miR\n-lacZ) - (AAV-miR\n-RXRa) 2.8788 1.04 35.4 2.758 0.2547  D2-MSNs 280 (AAV-miR\n-lacZ) - (AAV-miR\n-RXRa) 9.4070 1.10 52.4 8.543 <.0001  D1-MSNs 300 (AAV-miR\n-lacZ) - (AAV-miR\n-RXRa) 2.9913 1.04 35.4 2.866 0.2003  D2-MSNs 300 (AAV-miR\n-lacZ) - (AAV-miR\n-RXRa) 9.4070 1.10 52.4 8.543 <.0001  Results are averaged over some or all of the levels of: Sex  Degrees-of-freedom method: kenward-roger  P value adjustment: sidak method for 32 tests |
| Fig. 4h | Call : lmerTest ::lmer(sEPSC.freq ~ Virus*Sex*Cell.type + (1\|Mouse), control=lmerControl(check.nobs.vs.nRE="ignore", calc.derivs = F)) %>% anova(ddf = "Kenward-Roger")  Type III Analysis of Variance Table with Kenward-Roger's method  Sum Sq Mean Sq NumDF DenDF F value Pr(>F)  Virus 15.4868 15.4868 1 5.220 16.8442 0.008509 **  Sex 1.2372 1.2372 1 5.790 1.3457 0.291631  Cell.type 5.2797 5.2797 1 57.144 5.7425 0.019854 *  Virus:Sex 0.0788 0.0788 1 5.790 0.0857 0.779879  Virus:Cell.type 1.5555 1.5555 1 57.144 1.6919 0.198575  Sex:Cell.type 3.1076 3.1076 1 56.894 3.3800 0.071213 .  Virus:Sex:Cell.type 1.7459 1.7459 1 56.894 1.8989 0.173600  ---  Signif. codes: 0 ‘***’ 0.001 ‘**’ 0.01 ‘*’ 0.05 ‘.’ 0.1 ‘ ’ 1  Posthoc: emmeans(model, c(“Virus”, “Cell.type”)) %>% pairs(simple=”each”, adjust = “sidak”)  Cell.type Virus contrast estimate SE df t.ratio p.value  D1-MSNs . (AAV-miR\n-lacZ) - (AAV-miR\n-RXRa) 0.990 0.393 10.6 2.522 0.1117  D2-MSNs . (AAV-miR\n-lacZ) - (AAV-miR\n-RXRa) 1.615 0.403 14.1 4.004 0.0051  . AAV-miR\n-lacZ (D1-MSNs) - (D2-MSNs) 0.263 0.332 54.4 0.791 0.8960  . AAV-miR\n-RXRa (D1-MSNs) - (D2-MSNs) 0.888 0.347 59.0 2.561 0.0510  Results are averaged over some or all of the levels of: Sex  Degrees-of-freedom method: kenward-roger  P value adjustment: sidak method for 4 tests |
| Fig. 4i | Call : lmerTest ::lmer(sEPSC.freq ~ Virus*Sex*Cell.type + (1\|Mouse), control=lmerControl(check.nobs.vs.nRE="ignore", calc.derivs = F)) %>% anova(ddf = "Kenward-Roger")  Type III Analysis of Variance Table with Kenward-Roger's method  Sum Sq Mean Sq NumDF DenDF F value Pr(>F)  Virus 7.144 7.144 1 5.259 0.5445 0.492179  Sex 128.728 128.728 1 5.622 9.8107 0.022106 *  Cell.type 34.313 34.313 1 56.191 2.6151 0.111451  Virus:Sex 12.278 12.278 1 5.622 0.9357 0.373124  Virus:Cell.type 4.217 4.217 1 56.191 0.3214 0.573035  Sex:Cell.type 106.547 106.547 1 56.296 8.1202 0.006105 **  Virus:Sex:Cell.type 15.215 15.215 1 56.296 1.1596 0.286147  ---  Signif. codes: 0 ‘***’ 0.001 ‘**’ 0.01 ‘*’ 0.05 ‘.’ 0.1 ‘ ’ 1  Posthoc: emmeans(model, c(“Virus”, “Cell.type”)) %>% pairs(simple=”each”, adjust = “sidak”)  Cell.type Virus contrast estimate SE df t.ratio p.value  D1-MSNs . (AAV-miR\n-lacZ) - (AAV-miR\n-RXRa) -0.186 1.27 12.5 -0.147 0.9998  D2-MSNs . (AAV-miR\n-lacZ) - (AAV-miR\n-RXRa) -1.231 1.39 21.9 -0.885 0.8576  . AAV-miR\n-lacZ (D1-MSNs) - (D2-MSNs) -0.967 1.26 53.0 -0.770 0.9050  . AAV-miR\n-RXRa (D1-MSNs) - (D2-MSNs) -2.012 1.35 57.8 -1.493 0.4550  Results are averaged over some or all of the levels of: Sex  Degrees-of-freedom method: kenward-roger  P value adjustment: sidak method for 4 tests |
| Fig. 5a | Call: lmerTest ::lmer(Score ~ Group*Sex*PrePost + (1\|Mouse), control=lmerControl(check.nobs.vs.nRE="ignore", calc.derivs = F)) %>% anova(ddf = "Kenward-Roger")  Type III Analysis of Variance Table with Kenward-Roger's method  Sum Sq Mean Sq NumDF DenDF F value Pr(>F)  PrevPost 241469 241469 1 44 20.7223 0.00004165 ***  Group 58622 58622 1 44 5.0308 0.029986 *  Sex 14074 14074 1 44 1.2078 0.277747  PrevPost:Group 118341 118341 1 44 10.1557 0.002647 **  PrevPost:Sex 8768 8768 1 44 0.7524 0.390410  Group:Sex 4144 4144 1 44 0.3556 0.554009  PrevPost:Group:Sex 6939 6939 1 44 0.5955 0.444433  ---  Signif. codes: 0 ‘***’ 0.001 ‘**’ 0.01 ‘*’ 0.05 ‘.’ 0.1 ‘ ’ 1  Posthoc: emmeans(model, c(“Group”, “PrePost”)) %>% pairs(simple=”each”, adjust = “sidak”)  Group PrevPost contrast estimate SE df t.ratio p.value  HSV-GFP . (Pre-test) - (Post-test) -30.09 31.2 44.0 -0.965 0.8098  HSV-RXRa . (Pre-test) - (Post-test) -170.53 31.2 44.0 -5.472 <.0001  . Pre-test (HSV-GFP) - (HSV-RXRa) 1.19 37.9 79.7 0.031 1.0000  . Post-test (HSV-GFP) - (HSV-RXRa) -139.25 37.9 79.7 -3.679 0.0017  Results are averaged over some or all of the levels of: Sex  Degrees-of-freedom method: kenward-roger  P value adjustment: sidak method for 4 tests |
| Fig. 5b | Call: lmerTest ::lmer(Score ~ Group*Sex*PrePost + (1\|Mouse), control=lmerControl(check.nobs.vs.nRE="ignore", calc.derivs = F)) %>% anova(ddf = "Kenward-Roger")  Type III Analysis of Variance Table with Kenward-Roger's method  Sum Sq Mean Sq NumDF DenDF F value Pr(>F)  PrevPost 1296268 1296268 1 43 63.5737 0.0000000005172 ***  Group 123539 123539 1 43 6.0588 0.017925 *  Sex 484 484 1 43 0.0237 0.878326  PrevPost:Group 192428 192428 1 43 9.4374 0.003681 **  PrevPost:Sex 2680 2680 1 43 0.1314 0.718714  Group:Sex 1516 1516 1 43 0.0743 0.786418  PrevPost:Group:Sex 819 819 1 43 0.0401 0.842139  ---  Signif. codes: 0 ‘***’ 0.001 ‘**’ 0.01 ‘*’ 0.05 ‘.’ 0.1 ‘ ’ 1  Posthoc: emmeans(model, c(“Group”, “PrePost”)) %>% pairs(simple=”each”, adjust = “sidak”)  Group PrevPost contrast estimate SE df t.ratio p.value  HSV-GFP . (Pre-test) - (Post-test) -144.5 41.2 43.0 -3.505 0.0043  HSV-RXRa . (Pre-test) - (Post-test) -325.6 42.1 43.0 -7.725 <.0001  . Pre-test (HSV-GFP) - (HSV-RXRa) -20.8 54.0 73.9 -0.385 0.9920  . Post-test (HSV-GFP) - (HSV-RXRa) -201.9 54.0 73.9 -3.739 0.0014  Results are averaged over some or all of the levels of: Sex  Degrees-of-freedom method: kenward-roger  P value adjustment: sidak method for 4 tests |
| Fig. 5c | Call: lmerTest ::lmer(Score ~ Group*Sex*PrePost + (1\|Mouse), control=lmerControl(check.nobs.vs.nRE="ignore", calc.derivs = F)) %>% anova(ddf = "Kenward-Roger")  Type III Analysis of Variance Table with Kenward-Roger's method  Sum Sq Mean Sq NumDF DenDF F value Pr(>F)  PrevPost 547227 547227 1 41 33.8335 0.0000007886 ***  Group 107060 107060 1 41 6.6193 0.01381 *  Sex 16259 16259 1 41 1.0052 0.32193  PrevPost:Group 82817 82817 1 41 5.1204 0.02901 *  PrevPost:Sex 103312 103312 1 41 6.3875 0.01544 *  Group:Sex 234 234 1 41 0.0145 0.90476  PrevPost:Group:Sex 2547 2547 1 41 0.1575 0.69357  ---  Signif. codes: 0 ‘***’ 0.001 ‘**’ 0.01 ‘*’ 0.05 ‘.’ 0.1 ‘ ’ 1  Posthoc: emmeans(model, c(“Group”, “PrePost”)) %>% pairs(simple=”each”, adjust = “sidak”)  Group PrevPost contrast estimate SE df t.ratio p.value  AAV-miR-lacZ . (Pre-test) - (Post-test) -217.6 36.8 41.0 -5.906 <.0001  AAV-miR-RXRa . (Pre-test) - (Post-test) -95.7 39.3 41.0 -2.436 0.0750  . Pre-test (AAV-miR-lacZ) - (AAV-miR-RXRa) 61.5 54.7 64.8 1.125 0.7079  . Post-test (AAV-miR-lacZ) - (AAV-miR-RXRa) 183.4 54.7 64.8 3.354 0.0053  Results are averaged over some or all of the levels of: Sex  Degrees-of-freedom method: kenward-roger  P value adjustment: sidak method for 4 tests |
| Fig. 5d | Call: lmerTest ::lmer(Score ~ Group*Sex*PrePost + (1\|Mouse), control=lmerControl(check.nobs.vs.nRE="ignore", calc.derivs = F)) %>% anova(ddf = "Kenward-Roger")  Type III Analysis of Variance Table with Kenward-Roger's method  Sum Sq Mean Sq NumDF DenDF F value Pr(>F)  PrevPost 904021 904021 1 44 34.5725 0.0000005067 ***  Group 380007 380007 1 44 14.5326 0.000425 ***  Sex 622 622 1 44 0.0238 0.878130  PrevPost:Group 743787 743787 1 44 28.4447 0.0000031841 ***  PrevPost:Sex 5371 5371 1 44 0.2054 0.652623  Group:Sex 1414 1414 1 44 0.0541 0.817193  PrevPost:Group:Sex 4415 4415 1 44 0.1688 0.683141  ---  Signif. codes: 0 ‘***’ 0.001 ‘**’ 0.01 ‘*’ 0.05 ‘.’ 0.1 ‘ ’ 1  Posthoc: emmeans(model, c(“Group”, “PrePost”)) %>% pairs(simple=”each”, adjust = “sidak”)  Group PrevPost contrast estimate SE df t.ratio p.value  AAV-DIO-GFP . (Pre-test) - (Post-test) -18.04 46.7 44.0 -0.386 0.9920  AAV-DIO-RXRa . (Pre-test) - (Post-test) -370.12 46.7 44.0 -7.929 <.0001  . Pre-test (AAV-DIO-GFP) - (AAV-DIO-RXRa) 5.05 55.7 80.8 0.091 1.0000  . Post-test (AAV-DIO-GFP) - (AAV-DIO-RXRa) -347.03 55.7 80.8 -6.232 <.0001  Results are averaged over some or all of the levels of: Sex  Degrees-of-freedom method: kenward-roger  P value adjustment: sidak method for 4 tests |
| Fig. 5e | Call: lmerTest ::lmer(Score ~ Group*Sex*PrePost + (1\|Mouse), control=lmerControl(check.nobs.vs.nRE="ignore", calc.derivs = F)) %>% anova(ddf = "Kenward-Roger")  Type III Analysis of Variance Table with Kenward-Roger's method  Sum Sq Mean Sq NumDF DenDF F value Pr(>F)  PrevPost 140919 140919 1 44 5.3656 0.02526 *  Group 1 1 1 44 0.0000 0.99628  Sex 439 439 1 44 0.0167 0.89771  PrevPost:Group 2136 2136 1 44 0.0813 0.77682  PrevPost:Sex 102756 102756 1 44 3.9125 0.05421 .  Group:Sex 119 119 1 44 0.0045 0.94667  PrevPost:Group:Sex 14417 14417 1 44 0.5489 0.46269  ---  Signif. codes: 0 ‘***’ 0.001 ‘**’ 0.01 ‘*’ 0.05 ‘.’ 0.1 ‘ ’ 1  Posthoc: emmeans(model, c(“Group”, “PrePost”)) %>% pairs(simple=”each”, adjust = “sidak”)  Group PrevPost contrast estimate SE df t.ratio p.value  AAV-DIO-GFP . (Pre-test) - (Post-test) -86.06 46.8 44.0 -1.840 0.2602  AAV-DIO-RXRa . (Pre-test) - (Post-test) -67.19 46.8 44.0 -1.436 0.4974  . Pre-test (AAV-DIO-GFP) - (AAV-DIO-RXRa) -9.15 68.5 68.5 -0.134 0.9999  . Post-test (AAV-DIO-GFP) - (AAV-DIO-RXRa) 9.72 68.5 68.5 0.142 0.9998  Results are averaged over some or all of the levels of: Sex  Degrees-of-freedom method: kenward-roger  P value adjustment: sidak method for 4 tests |
| Fig. 5f | Call: lmerTest ::lmer(Score ~ Group*Sex*PrePost + (1\|Mouse), control=lmerControl(check.nobs.vs.nRE="ignore", calc.derivs = F)) %>% anova(ddf = "Kenward-Roger")  Type III Analysis of Variance Table with Kenward-Roger's method  Sum Sq Mean Sq NumDF DenDF F value Pr(>F)  PrevPost 484088 484088 1 44 22.1913 0.00002491 ***  Group 167540 167540 1 44 7.6803 0.0081505 **  Sex 28129 28129 1 44 1.2895 0.2622934  PrevPost:Group 293412 293412 1 44 13.4504 0.0006572 ***  PrevPost:Sex 53635 53635 1 44 2.4587 0.1240400  Group:Sex 35994 35994 1 44 1.6500 0.2056805  PrevPost:Group:Sex 21051 21051 1 44 0.9650 0.3312968  ---  Signif. codes: 0 ‘***’ 0.001 ‘**’ 0.01 ‘*’ 0.05 ‘.’ 0.1 ‘ ’ 1  Posthoc: emmeans(model, c(“Group”, “PrePost”)) %>% pairs(simple=”each”, adjust = “sidak”)  Group PrevPost contrast estimate SE df t.ratio p.value  Vehicle . (Pre-test) - (Post-test) -252.59 42.6 44.0 -5.924 <.0001  HX531 . (Pre-test) - (Post-test) -31.45 42.6 44.0 -0.738 0.9178  . Pre-test Vehicle - HX531 6.74 52.0 79.5 0.130 0.9999  . Post-test Vehicle - HX531 227.88 52.0 79.5 4.385 0.0001  Results are averaged over some or all of the levels of: Sex  Degrees-of-freedom method: kenward-roger  P value adjustment: sidak method for 4 tests |
| Fig. 6b | Call: stats::t.test(RXRa, GFP, var.equal = F)  Welch Two Sample t-test  data: RXRa and GFP  t = 2.3018, df = 8.5723, p-value = 0.04824  alternative hypothesis: true difference in means is not equal to 0  95 percent confidence interval:  0.1019231 20.8807635  sample estimates:  mean of x mean of y  74.76061 64.26926 |
| Fig. 6c | Call: lmerTest::lmer(resp ~ Group*Bin + (1\|RatID), control=lmerControl(check.nobs.vs.nRE="ignore", calc.derivs = F)) %>% anova(ddf = "Kenward-Roger")  Type III Analysis of Variance Table with Kenward-Roger's method  Sum Sq Mean Sq NumDF DenDF F value Pr(>F)  Bin 45462 5051.3 9 126 16.1586 < 0.00000000000000022 ***  Group 1314 1313.6 1 14 4.2020 0.05959 .  Bin:Group 15337 1704.1 9 126 5.4513 0.00000261 ***  ---  Signif. codes: 0 ‘***’ 0.001 ‘**’ 0.01 ‘*’ 0.05 ‘.’ 0.1 ‘ ’ 1  Posthoc: emmeans(model, c(“Group”)) %>% pairs(simple=”each”, adjust = “sidak”)  Bin contrast estimate SE df t.ratio p.value  Bin2 (HSV-GFP) - (HSV-RXRa) 0.000 10.8 71.6 0.000 1.0000  Bin3 (HSV-GFP) - (HSV-RXRa) 0.438 10.8 71.6 0.041 1.0000  Bin4 (HSV-GFP) - (HSV-RXRa) -0.438 10.8 71.6 -0.041 1.0000  Bin5 (HSV-GFP) - (HSV-RXRa) -0.375 10.8 71.6 -0.035 1.0000  Bin6 (HSV-GFP) - (HSV-RXRa) -0.688 10.8 71.6 -0.064 1.0000  Bin7 (HSV-GFP) - (HSV-RXRa) -1.438 10.8 71.6 -0.133 1.0000  Bin8 (HSV-GFP) - (HSV-RXRa) -10.062 10.8 71.6 -0.934 0.9872  Bin9 (HSV-GFP) - (HSV-RXRa) -26.438 10.8 71.6 -2.455 0.1534  Bin10 (HSV-GFP) - (HSV-RXRa) -51.062 10.8 71.6 -4.742 0.0001  Bin11 (HSV-GFP) - (HSV-RXRa) -48.375 10.8 71.6 -4.493 0.0003  Degrees-of-freedom method: kenward-roger  P value adjustment: sidak method for 10 tests |
| Fig. 6d | Call: lmerTest::lmer(intake ~ Group*Bin + (1\|RatID), control=lmerControl(check.nobs.vs.nRE="ignore", calc.derivs = F)) %>% anova(ddf = "Kenward-Roger")  Type III Analysis of Variance Table with Kenward-Roger's method  Sum Sq Mean Sq NumDF DenDF F value Pr(>F)  Bin 12.9718 1.44132 9 126 148.6600 <0.0000000000000002 ***  Group 0.0055 0.00550 1 14 0.5672 0.4639  Bin:Group 0.1055 0.01172 9 126 1.2085 0.2955  ---  Signif. codes: 0 ‘***’ 0.001 ‘**’ 0.01 ‘*’ 0.05 ‘.’ 0.1 ‘ ’ 1  Posthoc: emmeans(model, c(“Group”)) %>% pairs(simple=”each”, adjust = “sidak”)  Bin contrast estimate SE df t.ratio p.value  Bin2 (HSV-GFP) - (HSV-RXRa) 0.0000 0.0704 41.7 0.000 1.0000  Bin3 (HSV-GFP) - (HSV-RXRa) 0.0583 0.0704 41.7 0.828 0.9951  Bin4 (HSV-GFP) - (HSV-RXRa) -0.0329 0.0704 41.7 -0.467 1.0000  Bin5 (HSV-GFP) - (HSV-RXRa) -0.0155 0.0704 41.7 -0.220 1.0000  Bin6 (HSV-GFP) - (HSV-RXRa) -0.0163 0.0704 41.7 -0.232 1.0000  Bin7 (HSV-GFP) - (HSV-RXRa) -0.0192 0.0704 41.7 -0.272 1.0000  Bin8 (HSV-GFP) - (HSV-RXRa) -0.0751 0.0704 41.7 -1.067 0.9684  Bin9 (HSV-GFP) - (HSV-RXRa) -0.1093 0.0704 41.7 -1.552 0.7466  Bin10 (HSV-GFP) - (HSV-RXRa) -0.1225 0.0704 41.7 -1.740 0.6074  Bin11 (HSV-GFP) - (HSV-RXRa) -0.0645 0.0704 41.7 -0.916 0.9893  Degrees-of-freedom method: kenward-roger  P value adjustment: sidak method for 10 tests |
| Fig. 6f | Call: stats::t.test(RXRa, GFP, var.equal = F)  Welch Two Sample t-test  data: RXRa and GFP  t = 1.9504, df = 13.988, p-value = 0.07145  alternative hypothesis: true difference in means is not equal to 0  95 percent confidence interval:  -0.01428486 0.30073713  sample estimates:  mean of x mean of y  0.9174644 0.7742383 |
| Fig. 6g | Call: stats::t.test(RXRa, GFP, var.equal = F)  Welch Two Sample t-test  data: RXRa and GFP  t = 1.8907, df = 13.985, p-value = 0.07956  alternative hypothesis: true difference in means is not equal to 0  95 percent confidence interval:  -0.006179134 0.098068479  sample estimates:  mean of x mean of y  0.3022049 0.2562603 |
| Fig. 6h | Call: stats::t.test(RXRa, GFP, var.equal = F)  Welch Two Sample t-test  data: RXRa and GFP  t = 2.4123, df = 12.216, p-value = 0.03245  alternative hypothesis: true difference in means is not equal to 0  95 percent confidence interval:  0.00008343316 0.00160964925  sample estimates:  mean of x mean of y  0.001968249 0.001121708 |
| Fig. 6i | Call: stats::t.test(RXRa, GFP, var.equal = F)  Welch Two Sample t-test  data: RXRa and GFP  t = -2.3729, df = 8.0383, p-value = 0.0449  alternative hypothesis: true difference in means is not equal to 0  95 percent confidence interval:  -227.628605 -3.346908  sample estimates:  mean of x mean of y  112.4672 227.9549 |
| Fig. 6j | Call: stats::t.test(RXRa, GFP, var.equal = F)  Welch Two Sample t-test  data: RXRa and GFP  t = -3.2133, df = 8.6381, p-value = 0.01118  alternative hypothesis: true difference in means is not equal to 0  95 percent confidence interval:  -92.64515 -15.80614  sample estimates:  mean of x mean of y  41.49424 95.71988 |
| Fig. S1a | Call: voomlimma pairwise comparisons ^13^ Region \| Pairwise \| logFC \| AveExpr \| t \| P.Value \| adj.P.Val \| B; PfC \| CNvSN \| -0.0093388 \| 5.2569315 \| -0.1300432 \| 0.89718989 \| 0.98435258 \| -5.3125121; PfC \| CSvSS \| 0.00939849 \| 5.2569315 \| 0.11872913 \| 0.90608957 \| 0.99996252 \| -5.0458664; PfC \| SCvSS \| -0.0071641 \| 5.2569315 \| -0.0993019 \| 0.92139905 \| 0.99986625 \| -5.1044255; PfC \| CCvSS \| -0.010156 \| 5.2569315 \| -0.1408397 \| 0.88870985 \| 0.99998376 \| -5.1014521; NAc \| CNvSN \| 0.03696451 \| 5.69822834 \| 0.41106822 \| 0.68332724 \| 0.99988234 \| -5.0343219; NAc \| CSvSS \| -0.0981674 \| 5.69822834 \| -0.9116555 \| 0.36768669 \| 0.98608515 \| -4.7566445; NAc \| SCvSS \| -0.0291495 \| 5.69822834 \| -0.2720538 \| 0.78705062 \| 0.99999207 \| -4.9654173; NAc \| CCvSS \| 0.04840867 \| 5.69822834 \| 0.49075105 \| 0.62641938 \| 0.99967096 \| -4.960794; CPu \| CNvSN \| 0.00480925 \| 5.48510619 \| 0.08153178 \| 0.93540796 \| 0.9945433 \| -5.2007486; CPu \| CSvSS \| 0.01642379 \| 5.48510619 \| 0.25353179 \| 0.8010994 \| 0.9980441 \| -5.1172967; CPu \| SCvSS \| -0.0221368 \| 5.48510619 \| -0.3426836 \| 0.73355272 \| 0.99993911 \| -5.101094; CPu \| CCvSS \| 0.06589428 \| 5.48510619 \| 1.05860817 \| 0.29585374 \| 0.92441039 \| -4.7861407; BLA \| CNvSN \| -0.1022973 \| 5.86780444 \| -0.9520903 \| 0.34731392 \| 0.99995176 \| -4.7496382; BLA \| CSvSS \| -0.0221142 \| 5.86780444 \| -0.1956532 \| 0.84596678 \| 0.99985534 \| -4.9677128; BLA \| SCvSS \| -0.0046679 \| 5.86780444 \| -0.0452504 \| 0.96415439 \| 0.99377897 \| -5.0745682; BLA \| CCvSS \| -0.1485409 \| 5.86780444 \| -1.3812435 \| 0.17560234 \| 0.76010572 \| -4.4768464; vHipp \| CNvSN \| -0.0078934 \| 5.98335171 \| -0.098069 \| 0.92240992 \| 0.99941444 \| -4.9815669; vHipp \| CSvSS \| 0.02587779 \| 5.98335171 \| 0.2908456 \| 0.77280287 \| 0.90520304 \| -5.7349716; vHipp \| SCvSS \| -0.0405436 \| 5.98335171 \| -0.4739797 \| 0.63831553 \| 0.88467996 \| -5.5863417; vHipp \| CCvSS \| 0.11434033 \| 5.98335171 \| 1.22379411 \| 0.22881041 \| 0.87774886 \| -4.5628518; VTA \| CNvSN \| -0.0455038 \| 5.73956332 \| -0.6811418 \| 0.50104031 \| 0.99980889 \| -5.0900845; VTA \| CSvSS \| -0.038552 \| 5.73956332 \| -0.4702381 \| 0.64160792 \| 0.96114283 \| -5.0405085; VTA \| SCvSS \| -0.0057315 \| 5.73956332 \| -0.0822226 \| 0.93501944 \| 0.99785864 \| -5.2219874; VTA \| CCvSS \| -0.0780123 \| 5.73956332 \| -1.1135626 \| 0.27436748 \| 0.99963893 \| -4.775292 |
| Fig. S1b | Call: stats::cor.test(Rxra.rpkm, Addiction.Index, method = “pearson”) \| Cocaine \| cor \| All samples \| cor; PfC \| t = 1.0323, df = 17, p-value = 0.3164 \| 0.242883 \| t = 0.59009, df = 37, p-value = 0.5587 \| 0.09655691; NAc \| t = 3.7087, df = 18, p-value = 0.001607 \| 0.6581446 \| t = 3.0017, df = 36, p-value = 0.004855 \| 0.4474162; CPu \| t = 3.4438, df = 18, p-value = 0.002896 \| 0.6302283 \| t = 2.8951, df = 38, p-value = 0.006251 \| 0.4250977; BLA \| t = -0.43894, df = 14, p-value = 0.6674 \| -0.116514 \| t = -1.2284, df = 34, p-value = 0.2277 \| -0.2061486; vHipp \| t = 2.7544, df = 16, p-value = 0.01411 \| 0.5671446 \| t = 2.6527, df = 35, p-value = 0.01192 \| 0.409136; VTA \| t = -2.7199, df = 11, p-value = 0.01994 \| -0.634116 \| t = -2.772, df = 28, p-value = 0.009792 \| -0.4640448 |
| Fig. S2a | All statistics available in Table S1 |
| Fig. S2b | All statistics available in Table S2 |
| Fig. S2c | All statistics available in Table S3 |
| Fig. S2d | Call: stats::t.test(RXRa, GFP, var.equal = F)  Welch Two Sample t-test  data: RXRa and GFP  t = -7.3332, df = 4.4178, p-value = 0.00124  alternative hypothesis: true difference in means is not equal to 0  95 percent confidence interval:  -3.334636 -1.551632  sample estimates:  mean of x mean of y  1.000000 3.443134 |
| Fig. S2e | Call: stats::t.test(miR-RXRa, miR-lacZ, var.equal = F)  Welch Two Sample t-test  data: miR-RXRa and miR-lacZ  t = 8.4543, df = 22.58, p-value = 0.00000001899  alternative hypothesis: true difference in means is not equal to 0  95 percent confidence interval:  0.3703341 0.6106042  sample estimates:  mean of x mean of y  1.0000000 0.5095308 |
| Fig. S2f | Call : lm(norm.Relative.expression ~Virus*Cell.Type) %>% car::Anova(type = “III”, test.statistic = “F)  Anova Table (Type III tests)  Response: norm.Relative.expression  Sum Sq Df F value Pr(>F)  (Intercept) 23536.2 1 82.4719 0.00000001559 ***  Virus 22057.0 1 77.2889 0.00000002634 ***  Cell.Type 1361.7 1 4.7715 0.04101 *  Virus:Cell.Type 1317.9 1 4.6181 0.04406 *  Residuals 5707.7 20  ---  Signif. codes: 0 ‘***’ 0.001 ‘**’ 0.01 ‘*’ 0.05 ‘.’ 0.1 ‘ ’ 1  Posthoc: emmeans(model, c(“Virus”)) %>% pairs(simple=”each”, adjust = “sidak”)  contrast estimate SE df t.ratio p.value  (AAV-DIO-GFP) - (AAV-DIO-RXRa) -60.6 6.9 20 -8.791 <.0001  Results are averaged over some or all of the levels of: Cell.Type |
| Fig. S2g | Call: stats::t.test(RXRa, GFP, var.equal = F)  Welch Two Sample t-test  data: RXRa and GFP  t = 3.6574, df = 15.847, p-value = 0.002155  alternative hypothesis: true difference in means is not equal to 0  95 percent confidence interval:  0.1385732 0.5214286  sample estimates:  mean of x mean of y  1.330001 1.000000 |
| Fig. S2h | Call: stats::t.test(miR-RXRa, miR-lacZ, var.equal = F)  Welch Two Sample t-test  data: miR-RXRa and miR-lacZ  t = 5.2867, df = 13.186, p-value = 0.0001404  alternative hypothesis: true difference in means is not equal to 0  95 percent confidence interval:  0.2834615 0.6742673  sample estimates:  mean of x mean of y  1.0000000 0.5211356 |
| Fig. S2i | Call : lm(norm.Relative.expression ~Virus*Cell.Type) %>% car::Anova(type = “III”, test.statistic = “F)  Anova Table (Type III tests)  Response: norm.Relative.expression  Sum Sq Df F value Pr(>F)  (Intercept) 98.035 1 293.1128 0.0000000000002053 ***  Virus 25.023 1 74.8156 0.0000000341824499 ***  Cell.Type 0.012 1 0.0366 0.8503  Virus:Cell.Type 0.247 1 0.7373 0.4007  Residuals 6.689 20  ---  Signif. codes: 0 ‘***’ 0.001 ‘**’ 0.01 ‘*’ 0.05 ‘.’ 0.1 ‘ ’ 1  Posthoc: emmeans(model, c(“Virus”)) %>% pairs(simple=”each”, adjust = “sidak”)  contrast estimate SE df t.ratio p.value  (AAV-DIO-GFP) - (AAV-DIO-RXRa) -2.04 0.236 20 -8.650 <.0001  Results are averaged over some or all of the levels of: Cell.Type |
| Fig. S3b | Call: lmerTest::lmer(percent ~ Cell.type*Sex + (1\|mouseid) %>% anova(ddf = "Kenward-Roger")  Analysis of Variance Table  Response: percent  Df Sum Sq Mean Sq F value Pr(>F)  Sex 1 0.0 0.00 0.0000 1.0000  Test 3 5709.3 1903.09 291.6805 <2e-16 ***  Sex:Test 3 13.0 4.33 0.6639 0.5813  Residuals 28 182.7 6.52  ---  Signif. codes: 0 ‘***’ 0.001 ‘**’ 0.01 ‘*’ 0.05 ‘.’ 0.1 ‘ ’ 1  Posthoc: emmeans(model, c(“Cell.type”)) %>% pairs(simple=”each”, adjust = “sidak”)  contrast estimate SE df t.ratio p.value  (Drd2-pos Drd1-pos) - (Drd2-neg Drd1-pos) -29.22 1.21 28 -24.113 <.0001  (Drd2-pos Drd1-pos) - (Drd2-pos Drd1-neg) -26.68 1.21 28 -22.022 <.0001  (Drd2-pos Drd1-pos) - (Drd2-neg Drd1-neg) -30.86 1.21 28 -25.468 <.0001  (Drd2-neg Drd1-pos) - (Drd2-pos Drd1-neg) 2.53 1.21 28 2.091 0.2447  (Drd2-neg Drd1-pos) - (Drd2-neg Drd1-neg) -1.64 1.21 28 -1.355 0.7097  (Drd2-pos Drd1-neg) - (Drd2-neg Drd1-neg) -4.18 1.21 28 -3.446 0.0108  Results are averaged over some or all of the levels of: Sex  P value adjustment: sidak method for 6 tests |
| Fig. S3c | Call: lmerTest::lmer(Drd1 ~ Cell.type*Sex + (1\|mouseid) %>% anova()  Analysis of Variance Table  Type III Analysis of Variance Table with Satterthwaite's method  Sum Sq Mean Sq NumDF DenDF F value Pr(>F)  Sex 6.0827e+08 6.0827e+08 1 7.5 0.3036 0.5978  Test 9.8154e+13 3.2718e+13 3 19381.2 16327.8791 <2e-16 ***  Sex:Test 9.8641e+11 3.2880e+11 3 19381.2 164.0889 <2e-16 ***  ---  Signif. codes: 0 ‘***’ 0.001 ‘**’ 0.01 ‘*’ 0.05 ‘.’ 0.1 ‘ ’ 1  Posthoc: emmeans(model, c(“Cell.type”)) %>% pairs(simple=”each”, adjust = “sidak”)  contrast estimate SE df z.ratio p.value  (Drd2-pos Drd1-pos) - (Drd2-neg Drd1-pos) -36591 1919 Inf -19.067 <.0001  (Drd2-pos Drd1-pos) - (Drd2-pos Drd1-neg) 119496 1927 Inf 62.026 <.0001  (Drd2-pos Drd1-pos) - (Drd2-neg Drd1-neg) 118150 1918 Inf 61.604 <.0001  (Drd2-neg Drd1-pos) - (Drd2-pos Drd1-neg) 156087 834 Inf 187.199 <.0001  (Drd2-neg Drd1-pos) - (Drd2-neg Drd1-neg) 154741 810 Inf 190.949 <.0001  (Drd2-pos Drd1-neg) - (Drd2-neg Drd1-neg) -1346 824 Inf -1.633 0.4769  Results are averaged over some or all of the levels of: Sex  Degrees-of-freedom method: asymptotic  P value adjustment: sidak method for 6 tests |
| Fig. S3d | Call: lmerTest::lmer(Drd2 ~ Cell.type*Sex + (1\|mouseid) %>% anova()  Analysis of Variance Table  Type III Analysis of Variance Table with Satterthwaite's method  Sum Sq Mean Sq NumDF DenDF F value Pr(>F)  Sex 4.1526e+09 4.1526e+09 1 7.3 1.1311 0.3215  Test 2.3723e+14 7.9077e+13 3 19380.4 21539.9731 <2e-16 ***  Sex:Test 3.7516e+11 1.2505e+11 3 19380.4 34.0636 <2e-16 ***  ---  Signif. codes: 0 ‘***’ 0.001 ‘**’ 0.01 ‘*’ 0.05 ‘.’ 0.1 ‘ ’ 1  Posthoc: emmeans(model, c(“Cell.type”)) %>% pairs(simple=”each”, adjust = “sidak”)  contrast estimate SE df z.ratio p.value  (Drd2-pos Drd1-pos) - (Drd2-neg Drd1-pos) 184900 2598 Inf 71.183 <.0001  (Drd2-pos Drd1-pos) - (Drd2-pos Drd1-neg) -62384 2608 Inf -23.923 <.0001  (Drd2-pos Drd1-pos) - (Drd2-neg Drd1-neg) 181146 2596 Inf 69.779 <.0001  (Drd2-neg Drd1-pos) - (Drd2-pos Drd1-neg) -247284 1129 Inf -219.108 <.0001  (Drd2-neg Drd1-pos) - (Drd2-neg Drd1-neg) -3754 1097 Inf -3.423 0.0037  (Drd2-pos Drd1-neg) - (Drd2-neg Drd1-neg) 243529 1116 Inf 218.305 <.0001  Results are averaged over some or all of the levels of: Sex  Degrees-of-freedom method: asymptotic  P value adjustment: sidak method for 6 tests |
| Fig. S5b | Call: stats::t.test(RXRa, GFP, var.equal = F)  Welch Two Sample t-test  data: RXRa and GFP  t = 0.16909, df = 13.961, p-value = 0.8682  alternative hypothesis: true difference in means is not equal to 0  95 percent confidence interval:  -49.07437 57.47206  sample estimates:  mean of x mean of y  99.03560 94.83675 |
| Fig. S5c | Call: lmerTest::lmer(resp ~ Group*Bin + (1\|RatID), control=lmerControl(check.nobs.vs.nRE="ignore", calc.derivs = F)) %>% anova(ddf = "Kenward-Roger")  Type III Analysis of Variance Table with Kenward-Roger's method  Sum Sq Mean Sq NumDF DenDF F value Pr(>F)  Bin 46813 5201.4 9 126 29.3253 <0.0000000000000002 ***  Group 248 247.6 1 14 1.3959 0.2571  Bin:Group 1199 133.2 9 126 0.7509 0.6617  ---  Signif. codes: 0 ‘***’ 0.001 ‘**’ 0.01 ‘*’ 0.05 ‘.’ 0.1 ‘ ’ 1  Posthoc: emmeans(model, c(“Group”)) %>% pairs(simple=”each”, adjust = “sidak”)  Bin contrast estimate SE df t.ratio p.value  Bin2 (HSV-GFP) - (HSV-RXRa) 0.875 7.11 123 0.123 1.0000  Bin3 (HSV-GFP) - (HSV-RXRa) -0.250 7.11 123 -0.035 1.0000  Bin4 (HSV-GFP) - (HSV-RXRa) -0.125 7.11 123 -0.018 1.0000  Bin5 (HSV-GFP) - (HSV-RXRa) 2.250 7.11 123 0.317 1.0000  Bin6 (HSV-GFP) - (HSV-RXRa) -1.500 7.11 123 -0.211 1.0000  Bin7 (HSV-GFP) - (HSV-RXRa) -0.500 7.11 123 -0.070 1.0000  Bin8 (HSV-GFP) - (HSV-RXRa) -14.000 7.11 123 -1.970 0.4083  Bin9 (HSV-GFP) - (HSV-RXRa) -10.750 7.11 123 -1.512 0.7600  Bin10 (HSV-GFP) - (HSV-RXRa) -10.625 7.11 123 -1.495 0.7723  Bin11 (HSV-GFP) - (HSV-RXRa) -3.875 7.11 123 -0.545 0.9999  Degrees-of-freedom method: kenward-roger  P value adjustment: sidak method for 10 tests |
| Fig. S5d | Call: lmerTest::lmer(intake ~ Group*Bin + (1\|RatID), control=lmerControl(check.nobs.vs.nRE="ignore", calc.derivs = F)) %>% anova(ddf = "Kenward-Roger")  Type III Analysis of Variance Table with Kenward-Roger's method  Sum Sq Mean Sq NumDF DenDF F value Pr(>F)  Bin 14.8677 1.65196 9 126 51.4525 <0.0000000000000002 ***  Group 0.0005 0.00052 1 14 0.0163 0.9003  Bin:Group 0.2712 0.03014 9 126 0.9386 0.4943  ---  Signif. codes: 0 ‘***’ 0.001 ‘**’ 0.01 ‘*’ 0.05 ‘.’ 0.1 ‘ ’ 1  Posthoc: emmeans(model, c(“Group”)) %>% pairs(simple=”each”, adjust = “sidak”)  Bin contrast estimate SE df t.ratio p.value  Bin2 (HSV-GFP) - (HSV-RXRa) 0.20784 0.0896 140 2.320 0.1978  Bin3 (HSV-GFP) - (HSV-RXRa) -0.03333 0.0896 140 -0.372 1.0000  Bin4 (HSV-GFP) - (HSV-RXRa) -0.00940 0.0896 140 -0.105 1.0000  Bin5 (HSV-GFP) - (HSV-RXRa) 0.09298 0.0896 140 1.038 0.9722  Bin6 (HSV-GFP) - (HSV-RXRa) -0.03563 0.0896 140 -0.398 1.0000  Bin7 (HSV-GFP) - (HSV-RXRa) -0.00667 0.0896 140 -0.074 1.0000  Bin8 (HSV-GFP) - (HSV-RXRa) -0.10456 0.0896 140 -1.167 0.9400  Bin9 (HSV-GFP) - (HSV-RXRa) -0.04444 0.0896 140 -0.496 0.9999  Bin10 (HSV-GFP) - (HSV-RXRa) -0.02550 0.0896 140 -0.285 1.0000  Bin11 (HSV-GFP) - (HSV-RXRa) -0.00517 0.0896 140 -0.058 1.0000  Degrees-of-freedom method: kenward-roger  P value adjustment: sidak method for 10 tests |
| Fig. S5f | Call: stats::t.test(RXRa, GFP, var.equal = F)  Welch Two Sample t-test  data: RXRa and GFP  t = 0.12951, df = 11.097, p-value = 0.8993  alternative hypothesis: true difference in means is not equal to 0  95 percent confidence interval:  -0.1364311 0.1535098  sample estimates:  mean of x mean of y  0.8333136 0.8247742 |
| Fig. S5g | Call: stats::t.test(RXRa, GFP, var.equal = F)  Welch Two Sample t-test  data: RXRa and GFP  t = 0.12078, df = 11.046, p-value = 0.906  alternative hypothesis: true difference in means is not equal to 0  95 percent confidence interval:  -0.04502521 0.05025657  sample estimates:  mean of x mean of y  0.2751685 0.2725528 |
| Fig. S5h | Call: stats::t.test(RXRa, GFP, var.equal = F)  Welch Two Sample t-test  data: RXRa and GFP  t = 0.59101, df = 13.428, p-value = 0.5643  alternative hypothesis: true difference in means is not equal to 0  95 percent confidence interval:  -0.0002804119 0.0004925551  sample estimates:  mean of x mean of y  0.001396857 0.001290786 |
| Fig. S5i | Call: stats::t.test(RXRa, GFP, var.equal = F)  Welch Two Sample t-test  data: RXRa and GFP  t = -0.86583, df = 12.516, p-value = 0.4029  alternative hypothesis: true difference in means is not equal to 0  95 percent confidence interval:  -59.90295 25.72151  sample estimates:  mean of x mean of y  146.7488 163.8396 |
| Fig. S5j | Call: stats::t.test(RXRa, GFP, var.equal = F)  Welch Two Sample t-test  data: RXRa and GFP  t = -0.72621, df = 12.594, p-value = 0.481  alternative hypothesis: true difference in means is not equal to 0  95 percent confidence interval:  -31.01250 15.44647  sample estimates:  mean of x mean of y  59.79245 67.57547 |
